# Bayesian model discovery for reverse-engineering biochemical networks from data

**DOI:** 10.1101/2023.09.15.557764

**Authors:** Andreas Christ Sølvsten Jørgensen, Marc Sturrock, Atiyo Ghosh, Vahid Shahrezaei

## Abstract

The reverse engineering of gene regulatory networks based on gene expression data is a challenging inference task. A related problem in computational systems biology lies in identifying signalling networks that perform particular functions, such as adaptation. Indeed, for many research questions, there is an ongoing search for efficient inference algorithms that can identify the simplest model among a larger set of related models. To this end, in this paper, we introduce SLInG, a Bayesian sparse likelihood-free inference method using Gibbs sampling. We demonstrate that SLInG can reverse engineer stochastic gene regulatory networks from single-cell data with high accuracy, outperforming state-of-the-art correlation-based methods. Furthermore, we show that SLInG can successfully identify signalling networks that execute adaptation. Sparse hierarchical Bayesian inference thus provides a versatile tool for model discovery in systems biology and beyond.

## 1 Introduction

Bayesian statistical methods have become commonplace in many scientific disciplines. One reason for the victory march of Bayesian statistics lies in the ongoing development of different simulation-based inference methods, such as Approximate Bayesian Computation [ABC 30, 39, 58, 63], through which the Bayesian approach has become applicable to models without access to a closed-form likelihood. In essence, by comparing experimental data to a large set of simulations, researchers can determine the ranges of parameter values that can recover observations. Across many fields, researchers thus draw on sampling algorithms, such as Monte Carlo Methods or grid-based techniques, to constrain model parameters based on simulations and real-world data [e.g. 1, 32, 51].

In statistical inference, in addition to having to constrain model parameter values, one might not *a priori* know what mechanisms to include in models or how to capture system dynamics correctly. This problem lies at the root of model selection tasks. To give an example, consider the case of fitting time series (or snapshot) data to ordinary or partial differential equations (ODEs/PDEs) [cf. 2, 28, 36, 59]. The aim of such a fit is to constrain the values of the model parameters involved in the ODEs/PDEs. However, consider a scenario in which one must also determine a suitable functional form of the ODEs/PDEs. In other words, not only are the model parameter values unknown, but the terms to include in the ODEs/PDEs are also unknown. This dual task is known in machine learning as model discovery or *equation discovery* [e.g. 5, 6, 17, 42, 52]. Equation discovery is, hence, a large-scale model selection task between closely related models. A predefined set of parameters describes the most general and complex model, while simpler models (so-called submodels) are constructed by setting a subset of these parameters to zero. Therefore, the task of equation discovery becomes closely related to sparse regression and variable selection [44, 45]. Here, the term sparsity refers to reducing the number of non-zero parameters.

This paper presents a versatile algorithm for likelihood-free sparse inference based on an ABC Gibbs algorithm by Turner & Van Zandt [64]. We refer to our algorithm for sparse inference as SLInG (Sparse Likelihood-free Inference using Gibbs). Our algorithm can perform equation discovery but is not limited to dealing with ODEs/PDEs. Indeed, SLInG can be applied to any simulation-based sparse inference task. Moreover, like the Bayesian LASSO (least absolute shrinkage and selection operator) by Park & Casella that uses a Gibbs sampling algorithm [3, 37, 45], the algorithm by Turner & Van Zandt and hence SLInG are hierarchical. This property allows SLInG to autonomously adjust the level of sparsity invoked for each parameter in a data-dependent fashion. SLInG is thus placed among other adaptive approaches, such as Sorted L-One Penalized Estimation (SLOPE) [cf. 4, 12, and references therein]. In contrast to SLOPE, however, our approach is not limited to linear models.

This paper presents the application of SLInG to three case studies. In Section 3.1, we address a linear model for diabetes progression [9, 45]. The model provides a simple and well-studied example, allowing us to highlight and test the basic assets of our sparse sampling algorithm.

In Section 3.2, we use our algorithm to identify protein signalling networks that achieve biochemical adaptation, i.e. we look for network topologies with certain response properties [20, 31, 35, 40, 47, 50, 53]. Many studies in systems biology focus on understanding and mapping topologies that execute particular biological functions, such as adaptation. The search for efficient algorithms to accomplish this inference task is thus ongoing [e.g. 11, 13, 14, 35, 55, 56].

Finally, in Section 3.3, we employ our algorithm to identify links in stochastic gene regulatory networks from (synthetic) data, i.e. we seek to infer the specific network topology that underlies the data. This reverse engineering task poses a challenging problem in systems biology. Moreover, many of the existing methods for performing this reverse engineering are not based on mechanistic models and merely rely on constructing a correlation network [57]. We show that our algorithm can compete with other algorithms specifically tailored with this inference task in mind [38, 41].

## 2 Methods

In this paper, we consider the following problem: given a set of observations and a model with *K* parameters ***θ*** = *(θ*_1_, *θ*_2_, …, *θ*_*k*_), discard superfluous parameters (parameters that can be set to zero) and compute confidence intervals for the remaining parameters. This approach identifies the simplest sub-model(s) that can explain the data.

Within Bayesian statistics, our goal of sparse inference, i.e. variable selection, can be achieved by introducing suitable sparsity-inducing priors [44]. We discuss these priors in detail in Section 2.1. The priors themselves involve free parameters (***ϵ***), which implies that the problem can be phrased as a hierarchical model, i.e. ***ϵ*** are hyper-parameters. While it is commonplace to use the same hyper-parameters for all parameters [e.g., in LASSO 62], we assign a new hyper-parameter *ϵ*_*k*_ to each parameter, θ_*k*_. A similar approach is taken in the adaptive LASSO and SLOPE as mentioned in the introduction [3, 12, 67]. The use of parameter-specific hyper-parameters allows us to handle parameters of different orders of magnitude and to sift out redundant parameters without skewing the posterior distributions of the non-zero parameters.

Our Gibbs sampling algorithm is based on a Metropolis-Hastings algorithm. Our implementation builds on the hierarchical Approximate Bayesian Computation (ABC) algorithm by Turner & Van Zandt [64] and is summarised in Algorithm 1 in Appendix A. Figure 1 offers a schematic overview of our approach.

**Figure 1:**
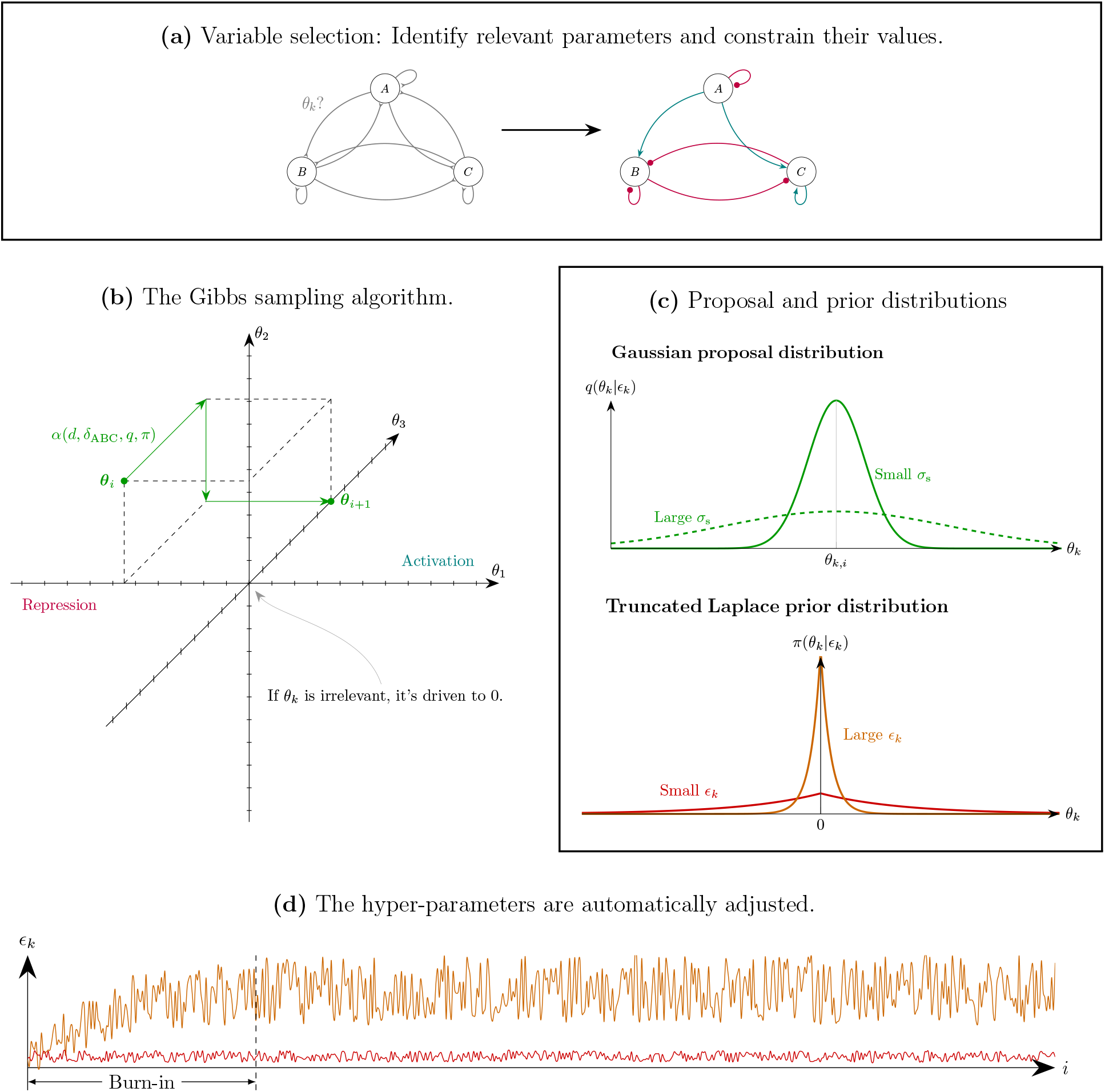
Schematic overview of the different components of SLInG. **(a)** Summary of sparse inference tasks exemplified through gene regulatory networks: We discard irrelevant links and constrain the properties of the remaining links. **(b)** To perform one step in the Markov chain starting from the parameter values at the ith iteration (***θ***_*i*_), the Gibbs sampler takes *K* perpendicular steps through the *K*-dimensional parameter space updating one parameter at a time. For each perpendicular step, one simulation is computed. The acceptance probability (*α)* at each perpendicular step depends on the parameter *δ*_ABC_ (that sets the overall level of sparsity), the proposal distribution *q*, the prior *π* (and hence the hyper-parameters, ***ϵ***), and the distance (*d*) between the predictions and the data. **(c)** The perpendicular steps are sampled from proposal distributions with adaptive width (*σ*_*s*_). Sparsity is imposed through parameter-specific priors. **(d)** The parameter-specific hyper-parameters (***ϵ***) of the priors are constantly updated.

To circumvent the intractable likelihood faced in many real-world applications, we substitute the likelihood in the acceptance rate of the Metropolis-Hastings algorithm with a kernel function *ψ*(*d*(*s*_sim_, *s*_obs_)/*δ*_ABC_) based on a distance measure *d*(*s*_sim_, *s*_obs_) between some summary statistics (*s*) of the simulation and the data. Here, *δ*_ABC_ denotes a tuning parameter. Also, note that it is the use of a distance measure between the summary statistics, d, that places the algorithm within the realm of ABC methods [58]. We discuss the distance measure, the summary statistics and the tuning parameter in detail in connection with each case study and 2.2. In this paper, we settle for the probability density function of the standard normal distribution as our kernel function.

The algorithm samples the parameter space in *N* steps going through all *K* parameters in each step. The hierarchical nature of the problem is included by treating model parameters and hyper-parameters in separate loops, sampling hyper-parameters based on parameter values from the previous sampling step. New values for *θ*_*k*_ are drawn from a proposal distribution *q*. Here, we use a normal distribution centred around the value of *θ*_*k*_ in the previous sampling step and with a variance of 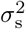 Like *δ*_ABC_, the user must choose 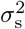. However, the chosen value of 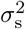 merely acts as an initial guess for a suitable step size. During the inference process, 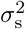 is constantly updated based on the variance of the samples in the Markov chain, i.e. unique values for 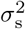 are automatically set for each parameter.

### 2.1 Sparsity inducing priors

Within the framework of Bayesian statistics, regression analysis employing the least absolute shrinkage and selection operator (LASSO) is imposed by introducing Laplace priors on the model parameters [e.g. 8, 45]. For a single parameter *θ*_*k*_, the Laplace distribution centred around zero is given by the probability density function

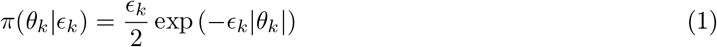

where *ϵ*_*k*_ is the corresponding hyper-parameter.

In addition, our implementation provides the option to define the region of the parameter space to explore. Our algorithm thus employs truncated sparsity priors by combining equation (1) with uniform priors. We use truncated Laplace priors for all case studies presented in this paper.

At the beginning of each new step in the Markov chain, we calibrate *ϵ*_*k*_ separately for each *θ*_*k*_. For this purpose, we consider a large number of uniformly distributed values of log_10_*(ϵ*_*k*_). For each of these values, we compute the probability of obtaining the current value of *θ*_*k*_ from the sparsity prior. We then randomly sample a new value of *ϵ*_*k*_ from the resulting distribution. In short, we thus sample *ϵ*_*k*_ from the conditional posterior based on the previous step (cf. Appendix A). In this manner, *ϵ*_*k*_ is iteratively adjusted: If the exploration of the parameter space yields a low value of *θ*_*k*_, a large value of *ϵ*_*k*_ will be imposed when using the Laplace prior, which further drives θ_*k*_ to zero unless the resulting changes in the distance measure provide evidence to the contrary. On the other hand, if the data points to a large value of *θ*_*k*_, *ϵ*_*k*_ will be small when using the Laplace prior, relaxing the imposed sparsity for this particular parameter.

### 2.2 Distance measures, tuning parameters and burn-in

The likelihood is often intractable when dealing with models that describe real-world dynamics. ABC methods, such as SLInG, circumvent this issue by introducing distance measures (*d*) that quantify the proximity between the data and the predictions of mathematical models [cf. 58]. For this purpose, ABC methods rely on suitable summary statistics **s** = (*s*_1_, *s*_2_, …, *s*_*N*_) that capture information in the simulated and observed data sets. What constitutes an appropriate distance measure is thus determined by the scientific question that one aims to address and by the data that one aims to fit. In this paper, we encounter three very distinct case studies, and we will define the appropriate distance metric in the corresponding subsections of the result section. To clarify the terminology, we note that the introduction of a distance measure means that ABC methods are said to be likelihood-free. However, when introducing the distance measure in the kernel of our Monte Carlo sampling scheme, we effectively introduce a surrogate likelihood [e.g. 51].

The sparsity of the fit is determined by *δ*_ABC_. If *δ*_ABC_ is too high, changes in d have little or no impact on the acceptance rate, i.e. the priors dominate the posterior, making the solution artificially sparse. On the other hand, if *δ*_ABC_ is too low, the Markov chain acceptance rate plummets, hindering convergence (see also Section 3.3). We thus explore, quantify and discuss the impact of our choice of *δ*_ABC_ for each case study.

In all case studies, we start our exploration of the parameter space from generic or randomised initial conditions. As a result, we need to discard a burn-in phase. During this burn-in phase, *δ*_ABC_ must be set to a sufficiently high value to allow for a proper exploration of the parameter space by smoothing out the impact that parameter changes have on the distance measure. Otherwise, the Gibbs sampler might get stuck at an excessively sparse solution. In other words, during the burn-in phase, *δ*_ABC_ might need to decrease to the desired value in a step-wise manner. Moreover, as a part of the burn-in, we introduce an initial grace period, during which the hyper-parameters are kept at low values, to allow a proper exploration of the parameter space.

Besides *δ*_ABC_, the width *(*σ**_*s*_) of the proposal distribution (q) determines the acceptance rate and sets the conditions for the exploration of the parameter space (cf. Appendix A). If the starting point of the Markov chain is far from the true parameter values, the proposal distribution should be broad. On the other hand, the parameter space around the best fit should be properly explored, requiring a narrow proposal distribution. Thus, *σ*_s_ must be iteratively adjusted during the run based on the previous samples.

Finally, to interpret the results obtained from the sampler, we need to make one more choice: If a parameter is superfluous, we obtain a narrow posterior distribution for the parameter around zero. Like for other sparse *Bayesian* inference methods, redundant parameters are not directly set to zero but take values close to zero. We thus need to choose, e.g., a threshold beyond which we interpret the parameter as being discarded by SLInG. Since this choice is problem-dependent, it is addressed separately for each individual application below (cf. Appendices B.1, B.3 and B.5).

## 3 Results

In this paper, we introduce SLInG for inferring simple (sparse) models describing a system using a Bayesian likelihood-free approach. In short, SLInG is a Gibbs sampling algorithm for ABC with sparsity-inducing priors that aims to minimise the distance (*d*) between the data and model predictions while discarding superfluous parameters. To achieve this goal, SLInG autonomously adjusts parameter-specific, sparsity-inducing hyper-parameters *(ϵ*_*k*_). The overall level of sparsity, i.e., the number of non-zero parameters, can be tuned by the parameter *δ*_ABC_. Intuitively, a smaller value of *δ*_ABC_ produces a closer fit to the data, which requires more complex (and less sparse) models. For further details, we refer the reader to Figure 1 and the methods section.

We present the applicability and performance of SLInG based on three case studies. First, we address a well-studied linear model to verify the performance of our approach in Section 3.1. Next, we identify biochemical signalling networks that possess certain input-output properties in Section 3.2. In this task, the sampler notably faces a non-linear model. Finally, turning our focus on a stochastic problem, we apply SLInG to learn gene regulatory networks from single-cell gene expression data in Section 3.3.

### 3.1 Linear models

The diabetes data set by Efron et al. [9] is often employed when demonstrating the performance of sparse sampling algorithms. The data set contains ten variables for 442 patients, and the task is to identify those parameters that are predictive of disease progression (see also Appendix B.1). The model that is assumed to describe the data set is a simple linear model [cf. 45].

To solve the task using SLInG, we employ the following distance measure:

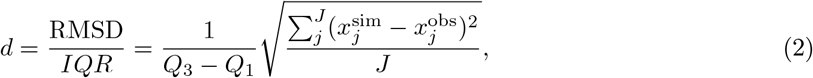

where 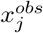 denotes the observed values of the response variable for the *J* patients and *Q*_1_ and *Q*_3_ denote the corresponding first and third quartiles (hence their difference is the interquartile range denoted as IQR). Here, 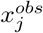 enters in the root mean square distance (RMSD) together with the corresponding prediction 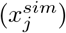 of the linear model — the model contains no intercept as both the features and the response variable are centred around their corresponding mean.

The results of our fit to the data are summarized in Fig 2. As expected, higher values of *δ*_ABC_ lead to sparser solutions. At intermediate values of *δ*_ABC_ (0.02 and 0.035), we find that important predictors of disease progression are BMI as well as two of the six serum measurements, including the glucose level, while lower values of *δ*_ABC_ indicate that other serum measurements and sex might play a role. This result is consistent with the literature [e.g. 45].

**Figure 2:**
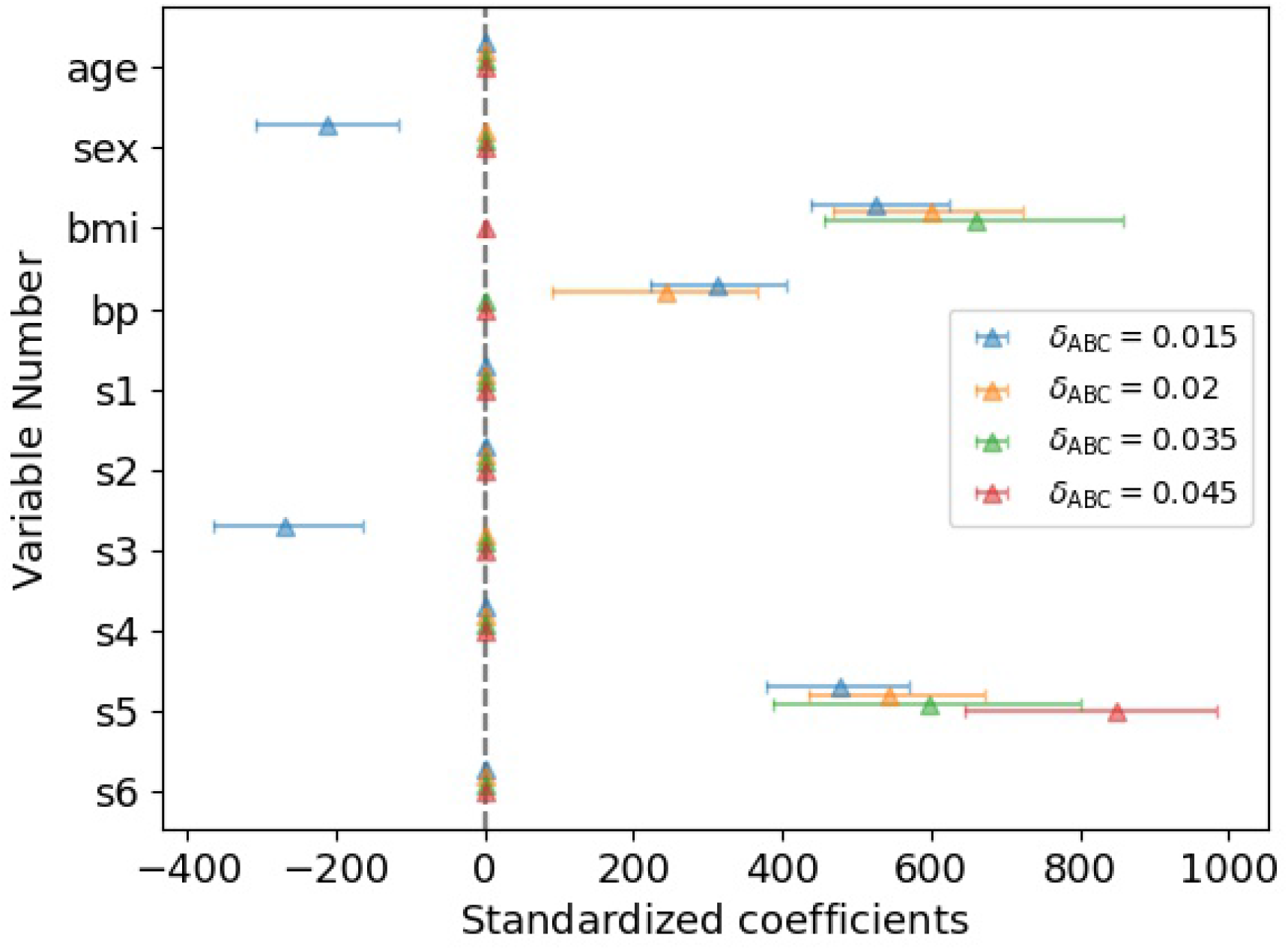
Parameter values inferred for the diabetes data set from Efron et al. [9] for different values of the overall sparsity parameter *δ*_ABC_. The plot shows the median (triangle) of each of the 10 parameters found by our sparse sampling algorithm. The error bars signify the corresponding 95 per cent confidence intervals. Each of the Markov chains that underlie the plot contains 8,500 samples after excluding a burn-in phase, in which *δ*_ABC_ is iteratively adjusted to the requested value.

To further quantify the performance of the Gibbs sampler, we used the data set from Efron et al. to construct synthetic data sets, i.e. we substituted the response variable with model predictions with known ground truth parameters and added noise (for details, see Appendix B.1). Since we know the parameter values that were used to create the synthetic data sets, this approach allows us to perform self-consistent tests. By doing so, we can check whether the Gibbs sampler is able to accurately reproduce the ground truth, i.e. the parameter values of the synthetic data. To map the impact of *δ*_ABC_, we repeat the inference for each of the synthetic data sets using different values of *δ*_ABC_. The results are summarized in Fig 3.

**Figure 3:**
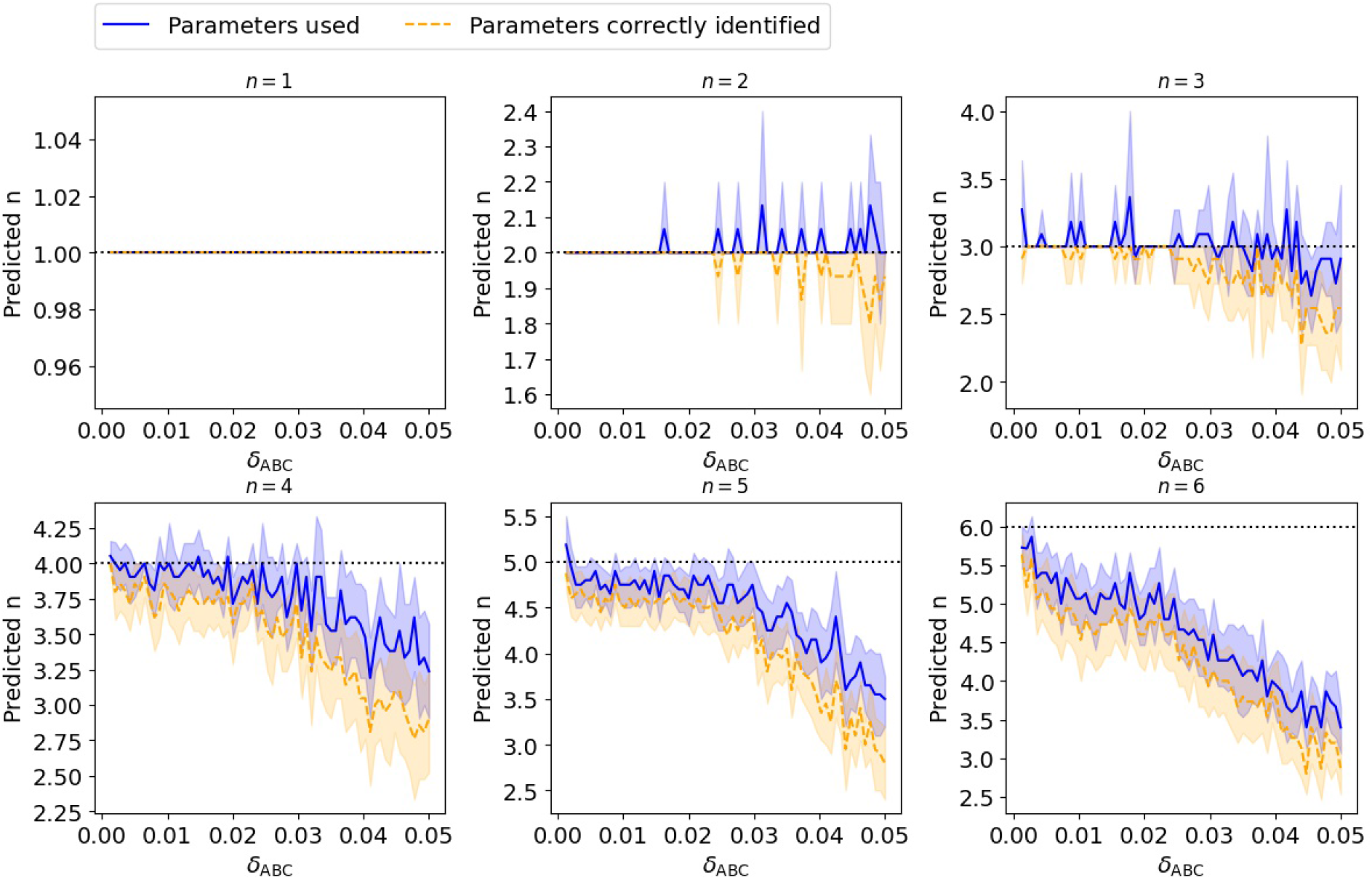
Parameters identified by SLInG as a function of *δ*_ABC_ for different values of the true number of parameters (*n*) employed to produce the synthetic data for our linear model. We fitted 100 synthetic data sets describing a simple linear system with 10 free parameters, varying the number of non-zero parameters (*n*) between 1 and 6 and the associated parameter values within two orders of magnitude. In each case, we repeated the inference for 66 values of *δ*_ABC_. The plot shows the mean number of non-zero parameters needed by the algorithm (blue) for each value of n and highlights the correctly identified parameters (orange) together with the corresponding 95 per cent confidence intervals. The plot is based on the best-fitting models from each Markov chain.

At low values of *δ*_ABC_, SLInG consistently recovers the ground truth, pinpointing the parameters that take non-zero values and discarding the rest. As *δ*_ABC_ increases, the sampler starts selecting models that are sparser than the ground truth. However, while the sampler points to solutions that are sparser than the model that generated the data, we note that the sampler points to parameters that are still a subset of the correct ones, i.e. the sampler picks up on the correct system dynamics. We also note that this behaviour is consistent with the results shown in Fig 2. For further details, we refer to Fig S1 in the supplementary material. Overall, we find that the performance of SLInG is not too sensitive to the value of our sparsity tuning parameter *δ*_ABC_, as long as this parameter is appropriately small.

While we only include the best-fitting models in Fig 3, we are not limited to providing a point estimate. Indeed, this is one of the great advantages of Bayesian approaches, such as Monte Carlo algorithms: SLInG provides the posterior distributions for the obtained parameter set. We illustrate this for one of our synthetic data sets in Fig 4. Also, the figure highlights that the algorithm autonomously sets the hyper-parameters (the *ϵ*_*k*_s) to be either very large (indicating that the corresponding parameter should be zero) or relatively small (suggesting the parameter of interest is set to have a non-zero value).

**Figure 4:**
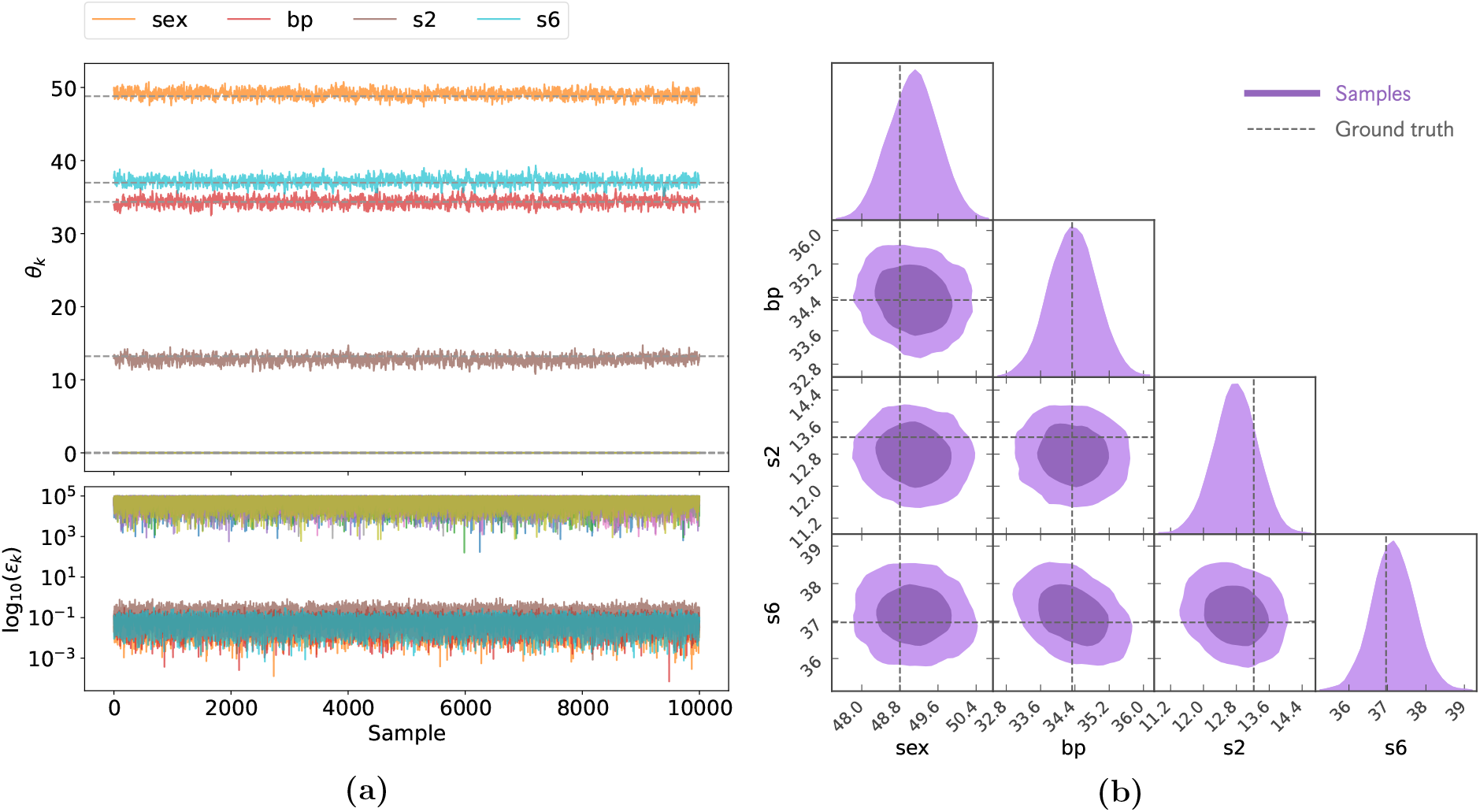
Markov chain for a single synthetic data set with four non-zero parameters after the exclusion of burn-in. (a) The upper panel shows the sampled parameter values (*θ*_*k*_) at each iteration, while the lower panel shows the corresponding hyper-parameters (*ϵ*_*k*_) in the Laplace prior. Note that we only include non-zero parameters in the legend while we plot all ten parameters and hyper-parameters across the 10,000 samples. In the upper panel, the dashed grey lines indicate the ground truth. (b) The resulting posterior distribution for the four non-zero parameters. All four parameters were correctly identified in this example.

### 3.2 Adaptation

Adaptive protein signalling networks respond to the changes in an input signal (*I*) with an output signal (*O*) (Figure 5a). Any change to the input signal (*I*_1_ → *I*_2_) initially leads to a strong response in the output signal (Figure 5b and c). However, after some time, the network will adapt to the change in the input signal, and the output signal will return close to its prestimulus level (Figure 5c). This kind of response occurs in different types of sensory systems, such as chemical receptors in olfactory cells [50]: While a new smell will initially provoke a strong response, it will disappear into the background with time. The chemotaxis of bacteria provides other examples of adaptive systems [10].

**Figure 5:**
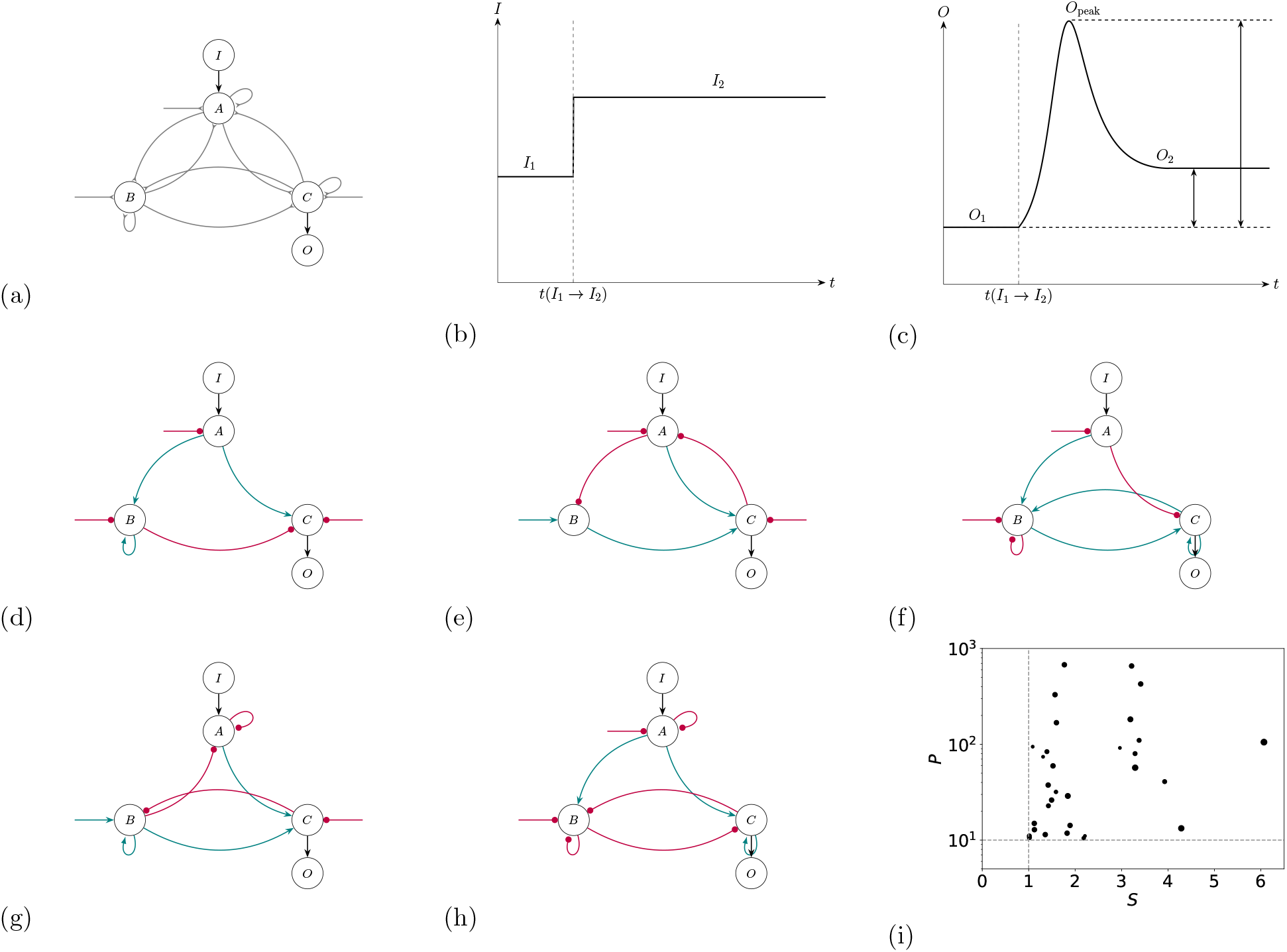
Summary figure for signalling networks exhibiting adaptation. (a) General protein network structure. Each network contains three proteins (A, B, C). (b) Change in the input (*I*). (c) Adaptive response in the output (*O*) to the change in the input. (d)-(h): Subsample of the adaptive networks found by the sampler. Red, blunt-headed arrows indicate repression, while cyan arrows indicate activation. Note, e.g., that (e) is an example of negative feedback, while (d) is an example of an incoherent feedforward loop. While we only show five networks here, our algorithm has successfully identified many more (cf. the supplementary material). (i) *S* and *P* for all unique networks listed in the supplementary material. The dashed line highlights the definition of adaptation: *S* > 1 and *P* > 10.

Ma et al. [35] set out to identify adaptive biochemical network topologies with three proteins: A, B, and C. A is receptive to the input, while C transmits the output. The three proteins might be linked to each other (thus repressing or activating each other) or possess self-feedback loops. Moreover, each protein might be activated or deactivated by other enzymes, for example, through phosphorylation/dephosphorylation reactions (*E*_*A*_, *E*_*B*_, *E*_*C*_). The supplementary material summarises the ordinary differential equations (ODEs) describing such networks (cf. Appendix B.2). In their paper, Ma et al. thus address the question of what particular network topologies result in adaptive behaviour. To answer this question, they define adaptation based on two quantities that they denote as sensitivity and precision. The sensitivity (*S*) encapsulates the initial reaction of the network:

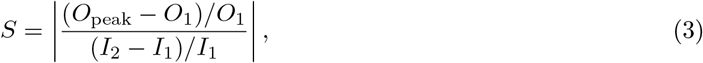

where *O*_1_ and *O*_peak_ denote the initial and extreme value in the output signal. *I*_1_ and *I*_2_ are the initial and final input signals. The precision (*P*) quantifies the successful return to the prestimulus level:

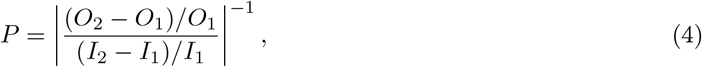

where *O*_2_ denotes the final output signal. Networks are said to be adaptive when both *S* > 1 and *P* > 10.

Ma et al. identify adaptive networks through a brute-force search of all network topologies in their parameter space. To address this rather abstract task with SLInG, we use the following distance measure:

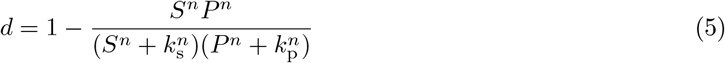

that is, a non-linear, increasing function of sensitivity and precision. We set *n* = 2, *k*_s_ = 1, and *k*_p_ = 10. Like Ma et al., we set *I*_1_ = 0.5 and *I*_2_ = 0.6. The priors of the model parameters and details of the set-up are described in Appendix B.2) and Appendix B.3.

Based on one hundred independent Markov chains, we identified 31 unique topologies showing adaptation. Figure 5 shows five examples. All networks are included in the supplementary material (Figs S2-S3). The 31 successful topologies were all found when choosing low values of *δ*_ABC_. This behaviour was to be expected as higher values of *δ*_ABC_ imply that the imposed sparsity requirements pull the sampler away from the relevant region of the parameter space.

Ma et al. suggest that two topologies lie at the heart of any adaptive network: a negative feedback loop with a buffer node (node B in Fig 5) and an incoherent feedforward loop with a proportioner (also node B in Fig 5). Indeed, all the networks identified by SLInG fall into these categories. In Figure 5, panel (e) is an example of a negative feedback loop: A activates C, but C represses A. Meanwhile, in Figure 5, panel (d) is an example of an incoherent feedforward loop: A activates C, but through its activation of B, A also represses C. In addition, several of the networks are found to use a combination of these strategies.

In this Section, our aim is to identify topologies that allow for adaptation. Our research question is thus about the existence of such topologies: We are interested in whether we can construct sparse networks that exhibit adaptation but not in the precise parameter values. Thus, in contrast to the analysis presented in Section 3.1, we do not aim to constrain the parameter values. Since we are not making any statements about the posterior distribution of the parameter values, we do not need to ensure convergence of the Markov chain. However, a closer look at the Markov chain might still give additional insights into the identified signalling network. As noted by Ma et al., one might ask whether any given network topology robustly results in adaptation for a broad range of parameter values or whether some parameters need to be fine-tuned to ensure adaptation. Each of the networks might thus either correspond to a broad or a narrow peak in the posterior probability landscape. Indeed, one of the advantages of using a Monte Carlo approach is that one can detect fine-tuned topologies that would be overlooked in a coarse grid search of the parameter space. Just like in the case of a grid search, we can make qualitative statements on whether the identified topologies robustly lead to adaptation if we have obtained enough independent samples — even if the Markov chain has not converged to the posterior. We illustrate this point in Figures S4 and S5 in the supplementary material. The network topology illustrated in Figure S4 is more robust than the one topology illustrated in Figure S5 as evident by the wider range of parameter values resulting in adaptation.

Importantly, our case study illustrates that SLInG can go beyond the simple linear models exemplified in Section 3.1. As discussed in the supplementary material, the ODEs used to model this system are relatively high-dimensional, including 26 free parameters, and are also non-linear.

### 3.3 Stochastic regulatory genetic networks

Having demonstrated that our sparse sampling algorithm can be successfully applied to linear and non-linear models in Sections 3.1 and 3.2, we proceed to discuss a stochastic example: The reverse engineering of gene regulatory networks from single-cell transcription data.

The topologies of gene regulatory networks can be inferred from single-cell gene expression data, e.g., single-cell RNA-sequencing data. However, this inference task is challenging because gene expression is known to be affected by intrinsic noise in biochemical reactions [54]. Intrinsic noise can thus lead to significant variability in gene expression levels, even for genes that are regulated by the same factors. This variability can obscure the true underlying regulatory relationships and make it challenging to distinguish real regulatory interactions from noise. In addition, typically, we do not have access to single-cell time-course data and/or perturbation data, so we have to only rely on snap-shot data.

To illustrate the performance of SLInG for model-based inference of genetic networks from data, we choose to work with synthetic data since this allows us to perform a self-consistent test and hence to benchmark SLInG (see also Section 3.3.1). Here, we focus on gene regulatory networks that contain four genes. The links between these genes might either lead to repression or activation of gene expression. We do not consider self-activation or self-repression. Gene-regulatory networks are often simulated using the Gillespie algorithm that respects the discreteness of the molecules [18]. However, in this paper, we employ a stochastic differential equations (SDE) approach due to its computational efficiency (see Appendix B.4). For this purpose, we employ a recently published library for biochemical modelling by Loman et al. [33] to produce synthetic data and to perform a self-consistent test, i.e. SLInG draws on the same library to infer the model parameters that underlie the synthetic data. The output of the model is a distribution of gene expression counts across single cells, capturing gene-gene correlations that contain information with regard to the underlying network (cf. Figure 6c below). The output of the simulations hereby closely mimics real-world experiments with cell populations. Here, we focus on the difficult case of working with only snap-shot data and do not include time-course data and/or perturbation data.

**Figure 6:**
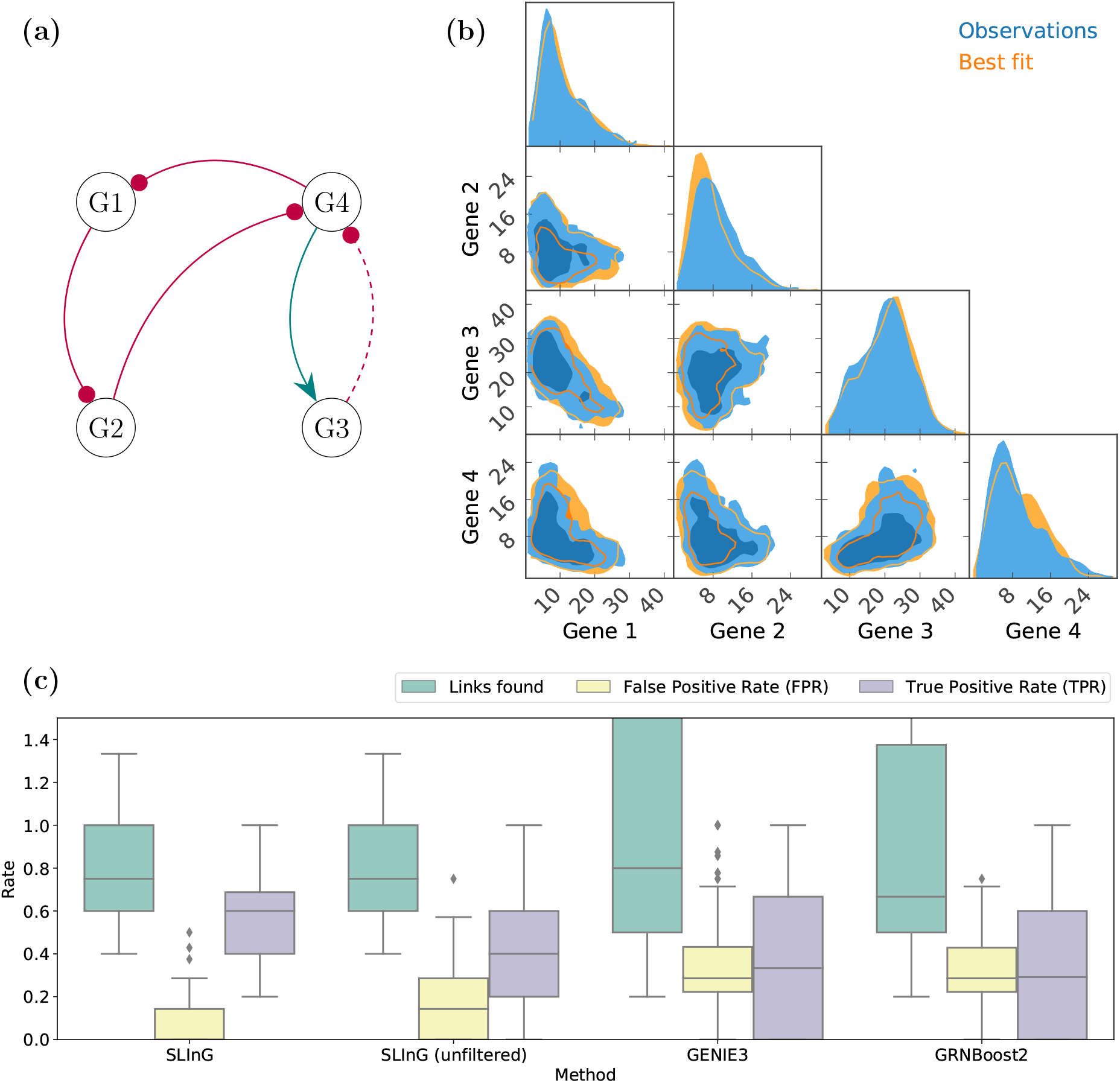
Performance summary for stochastic models of gene regulatory networks. (a) The network that underlies the particular synthetic data. The Gibbs sampler correctly identified four out of the five links — the link that was not found is indicated with a dashed line. (b) Example of predicted and observed gene expression across 1,000 cells. (c) We compare our sparse Gibbs sampling algorithm (SLInG) with two established gene network inference tools from the literature, GENIE3 and GRNBoost2. We include the results for SLInG for the 56 simulations that passed our quality check, as well as for all 100 networks. The plot includes the number of links that the algorithms settle for as a fraction of the true number of links. Moreover, we list the true and false positive rates, summarising how many links were correctly identified and how many spurious links were claimed to be found. The true and false positive rates are defined as *T PR* = *T P*/*P* and *FPR* = *FP*/*N*, respectively. *T P* and *FP* denote the total number of true and false positives, respectively, while *P* and *N* denote the numbers of positives and negatives in the data. Thus, while *T P* and *FP* stem from a comparison between the model predictions and the ground truth, *P* and *N* are based on the ground truth alone.

To fit the data, we use the first two moments of gene expression as summary statistics [66] and employ the following Euclidean distance:

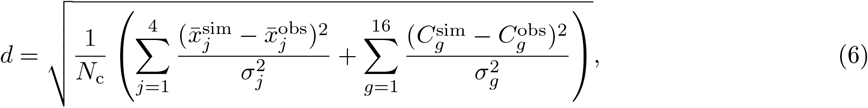

where 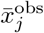 and 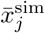 denote the observed and predicted mean expression of gene *j*, respectively. The standard deviation (*σ*_*j*_) of the observed mean expression is determined by bootstrapping. 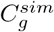 and 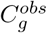 denote the 16 elements of the covariance matrices of the obtained distribution of gene expressions for the model predictions and the synthetic observations, respectively. The standard deviations (*σ*_*g*_) of the element of the covariance matrix of the observations are found by bootstrapping. Finally, *N*_*c*_ denotes the number of cells in the experiment. Here, the cell population is assumed to contain 1,000 cells. We created one hundred synthetic data sets based on random 4-node gene-regulatory networks. For further details, we refer the reader to Appendix B.5.

In Sections 3.1 and 3.2, we show that higher values of *δ*_ABC_ yield sparser solutions and that we fully employ the information in the data when setting *δ*_ABC_ to a very low value. In the present case, this approach fails. Since the model is stochastic, SLInG is prone to get stuck if it becomes too sensitive to changes in the distance measure — after all, the distance measure will vary between two samples with the *same* parameter values due to stochasticity and the finite sample size (1000 cells). Indeed, Monte Carlo methods are known to struggle with stochastic simulations [e.g. 24, 27, 65]. However, if we increase the value of *δ*_ABC_, the sampler becomes less sensitive to the stochasticity of the simulations and hence becomes effective for the inference based on stochastic simulations. Just by varying *δ*_ABC_, we can thus overcome an obstacle that would otherwise severely stifle our algorithm. For further details on the synthetic data and priors, we refer the reader to Appendix B.5.

We fitted each synthetic data set using SLInG and subsequently assessed the quality of each fit (cf. Appendix B.5). If we were faced with a low-quality fit when applying the algorithm to real-world data, we could try to improve the fit by, e.g., adjusting initial conditions. Here, however, we merely discard low-quality fits and note that the algorithm passed our quality check out of the box in 56 per cent of the cases.

On average, the number of non-zero links in the best-fitting models corresponds to 79 per cent of the number of links used to create the synthetic data set in question. We note that it is to be expected that the algorithm will not find all links due to the requirement of sparsity, our conservative cut-off in the link strength, and the expected trade-off between the link strengths, the values taken by base expression rates and model simplicity (Appendix B.4). Notably, 80 per cent of the predicted links correctly identify a link between genes in the synthetic data *and* attribute the correct sign to the interaction — that is, we strictly distinguish between repressive and activating links. In other words, 80 per cent of the predicted links are really present in the data. Combining these two numbers, we achieve a true positive rate of 60 per cent, i.e., on average, the algorithm correctly identified 60 per cent of all the links present in the data. Moreover, in every single case, at least one link was correctly identified. This is a very promising result. Indeed, as discussed in Section 3.3.1 and as shown in Fig 6c, our sampler outperforms algorithms explicitly designed to identify links in gene regulatory networks.

To illustrate the performance of SLInG further, we include an example of a fit to the synthetic gene expression data in Fig 6b.

#### 3.3.1 Comparison to other algorithms

We compare our method with two of the most commonly used and high-performing gene regulatory network inference algorithms: GENIE3 and GRNBoost2 [see 22, 26, 41, 61, as well as Appendix B.6 for further details]. Both algorithms are well-established in the field. By combining their output with the ‘ppcor’ package in R to compute the directionality of the interactions [25], we can produce estimates of signed, directed networks that we can compare with SLInG.

As can be seen from Fig 6c, both GENIE3 and GRNBoost2 often overestimate the true number of links, while the median number of links obtained with both algorithms is similar to that obtained with SLInG. Moreover, GENIE3 and GRNBoost2 lead to a significantly higher number of false positives and a somewhat lower number of true positives than SLInG. Even if we were to fix the number of links to the ground truth, this statement would still hold true. The high number of false positives is thus not solely an artefact that stems from overestimating the true number of links.

We note that SLInG has an advantage that further boosts its scores in our comparison: The output of SLInG allows us to compare the predicted network structure with the data directly. We can hence impose a quality check based on which we can discard bad fits. GENIE3 and GRNBoost2 allow for no such filtering. If we abandon the quality check offered by SLInG, its median true positive rate drops from 0.6 to 0.4, lying closer to that of both GENIE3 and GRNBoost2 (cf. Fig 6). This being said, one of the appeals of using SLInG lies in its transparency, and one wouldn’t abandon such quality checks when dealing with real-world data.

## 4 Discussion

Inference of genetic and signalling networks is a fundamental and challenging task in systems biology. Some of the current methods rely on obtaining correlation networks and are, therefore, not by nature mechanistic. Using ODEs or stochastic models of biochemical networks is challenging as the network inference task would be equivalent to large-scale model selection among many competing network topologies. In this paper, we cast the problem as model/equation discovery using Bayesian sparse inference and demonstrate that our implementation reliably pinpoints the correct non-zero parameters relating to the regulatory links existing in the network in self-consistent tests. Our approach relies on a sparsity-inducing Gibbs ABC algorithm that we call SLInG. It is applicable to any simulation-based inference task, including for non-linear or stochastic models. Furthermore, the algorithm successfully adjusts the level of sparsity in a data-dependent fashion for each parameter individually, resulting in models with the appropriate level of complexity.

In Section 3.1, we address a well-studied linear model for the occurrence of diabetes among patients. We demonstrate that SLInG correctly identifies the non-zero parameters when faced with synthetic data and yields results that are consistent with the literature when dealing with real-world data. We show that our Bayesian algorithm yields the joint posterior distribution of all parameters, quantifying the uncertainty of the model predictions [see also 17, 21, 45, for other Bayesian algorithms for equation discovery that provide posteriors]. In contrast, many sparse sampling schemes only provide point estimates — say, classical LASSO or some state-of-the-art approaches within equation discovery [e.g. 5, 6].

In Section 3.2, we successfully identify signalling networks that exhibit a particular biochemical function: Adaptation. Following Ma et al. [35], who performed an exhaustive grid search to identify such systems, we consider signalling networks with three genes. Such systems are well-studied, allowing for a comparison to the literature. We thus see that SLInG consistently points to signalling network topologies that show the expected features: negative feedback loops with a buffer node and incoherent feedforward loops with a proportioner. Moreover, based on the samples in the Markov chain, we can directly compare the robustness of different network topologies.

An exhaustive search of all network topologies and a grid parameter search in each case, such as the one performed by Ma et al., fully covers a region of the parameter space at a predetermined resolution, which might become computationally insurmountable as the number of parameters increases [31, 35, 53]. In contrast, a Monte Carlo algorithm, such as SLInG, accumulates more samples from regions with high posterior probability, efficiently combining scouting for topologies and inferring parameters that meet given selection criteria. It would thus be computationally feasible to extend the search for adaptive topologies beyond signalling networks with three genes using SLInG, while the same would not hold true for a grid search of the parameter space.

In this connection, we note that while our Gibbs sampling algorithm scales relatively well when dealing with higher dimensional problems, it is inevitable that we will face the curse of dimensionality for vastly more extensive networks. One problem is the computational cost of the increased number of simulations required. However, in the last few years, a promising new line of research based on different machine learning approaches has emerged that can accelerate simulation-based inference [34, 60], and some of these approaches can be applied in tandem with SLInG to overcome the stumbling block imposed by computational costs [24]. An interesting future direction would be adopting these methods for model discovery and sparse Bayesian inference [29].

Moreover, by looking for adaptive signalling networks, we demonstrate that SLInG can successfully handle parameters that do not simply enter as coefficients in a linear combination of source terms. In contrast, state-of-the-art methods in equation discovery, such as sparse identification of non-linear dynamics (SINDy), are designed to constrain a matrix of linear combination coefficients [**Ξ**, e.g. 5, 6, 17, 21]. In other words, while the resulting dynamics are thus non-linear, the equations explored by SINDy must take the form 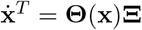 where **x** denotes the state of the system and **Θ**(**x**) denotes a matrix of library terms that may enter the ODEs that describe the systems dynamics.

As discussed above, some authors have performed exhaustive searches over the space of all network topologies to address this kind of non-linear model discovery problem in the context of systems biology. Other authors have used methods based on genetic algorithm and *in silica* evolution [11, 13, 14]. These methods draw on a library of genotypes that represent specific networks and their parameter sets. The algorithms use rounds of mutations and selection to find fitter networks. In principle, these methods could explore an unlimited level of complexity to obtain solutions given an appropriate set-up for the allowed mutations in the system. In contrast, in our approach, we specify the maximum level of complexity at the outset and look for the simplest sub-networks that perform the desired task or fit the available data. Our case study of adaptive systems suggests that SLInG can provide a more versatile and efficient option to the existing *in silica* evolution approaches.

In Section 3.3, we reverse engineer the topology of gene regulatory networks based on gene expression data. Like the identification of signalling networks with particular functions, inference of genetic networks from gene expression data is an important open problem in computational systems biology [57]. Many algorithms have thus been proposed to address this task and have, over the years, been applied to bulk, single-cell, temporal and perturbation data. Many of the existing methods are not based on mechanistic models, and they merely try to reconstruct a correlation network. We demonstrate that SLInG shows good performance when compared to some of the leading correlation-based algorithms (GENIE3 and GRNBoost2) that are specifically tailored to reverse engineer gene regulatory networks. Moreover, we note that we perform the reverse engineering of signalling networks based on stochastic models (SDEs). While other Monte Carlo algorithms struggle to explore the parameter space when faced with stochastic models, we show that our implementation does not suffer from this shortcoming.

Correlation-based inference methods for genetic networks, such as GENIE3 and GRNBoost2, are computationally efficient and scale well with the size of the networks. However, mechanistic models can systematically take the experimental design into account and combine multiple types of data such as single-cell, bulk, temporal and perturbation gene expression data. Indeed, a very promising area of future work is harnessing the information contained in the multi-modal single-cell data using inference based on mechanistic models [7, 19, 49].

When quantifying the performance of an inference framework, it is standard procedure to perform benchmarking based on synthetic data, i.e. to perform self-consistent tests, as this allows us to compare to the ground truth [e.g. 24, 34, 48]. Thus, to test SLInG while avoiding *systematic* errors that arise from model misspecification [43], we provide the sampler with the same modelling libraries that are used to generate the synthetic data in all our case studies. However, when interpreting these bench-marking results, one must bear in mind that a self-consistent test is an idealised situation and that all models are simple abstractions of real phenomena. Whether a good performance in self-consistent tests of simulation-based approaches translates into a good performance when dealing with real-world data, therefore, depends on how well the mathematical model captures the phenomenon in question. Moreover, the choice of distance measure can contribute to reducing the impact of model misspecification [15].

In sparse regression, setting the level of sparsity is a difficult task; methods such as cross-validation can be used, but they are very time-consuming [23]. To address this issue, SLInG entails hyper-parameters that adjust the sparsity-inducing prior for each parameter (similar to adaptive LASSO and SLOPE [3, 12, 67]) and a tuning parameter *(δ*_*ABC*_) that affects the quality of the fit to data and hence the overall level of sparsity. The decision about the level of sparsity is ultimately closely related to how much we believe in the data, as complex models will have a tendency to overfit the data. Similarly, the level of sparsity is related to how much we believe in our model due to the issue of model misspecification in real-world applications discussed above. In practice, we obtain reasonable results as long as we choose *δ*_*ABC*_ to be relatively small in our case studies. Also, the value of each hyper-parameter gives a clear indication of which parameters should be set identically to zero. Overall, this makes SLInG efficient and powerful in inferring models with the right level of complexity.

In this paper, we demonstrate that SLInG is a flexible framework that is well-equipped to perform sparse inference based on a wide variety of simulations, including non-linear and stochastic models. Not only does the presented hierarchical Bayesian sampling scheme thus provide an alternative to other algorithms, but it is also a versatile tool that can be used to address multiple research questions in systems biology and beyond.

## Acknowledgement

ACSJ is supported by the Eric and Wendy Schmidt AI in Science Postdoctoral Fellowship, a Schmidt Futures program. We acknowledge conversations with Philipp Thomas.

## A Algorithm overview

### Algorithm 1 Sparse likelihood-free inference using Gibbs (SLInG)

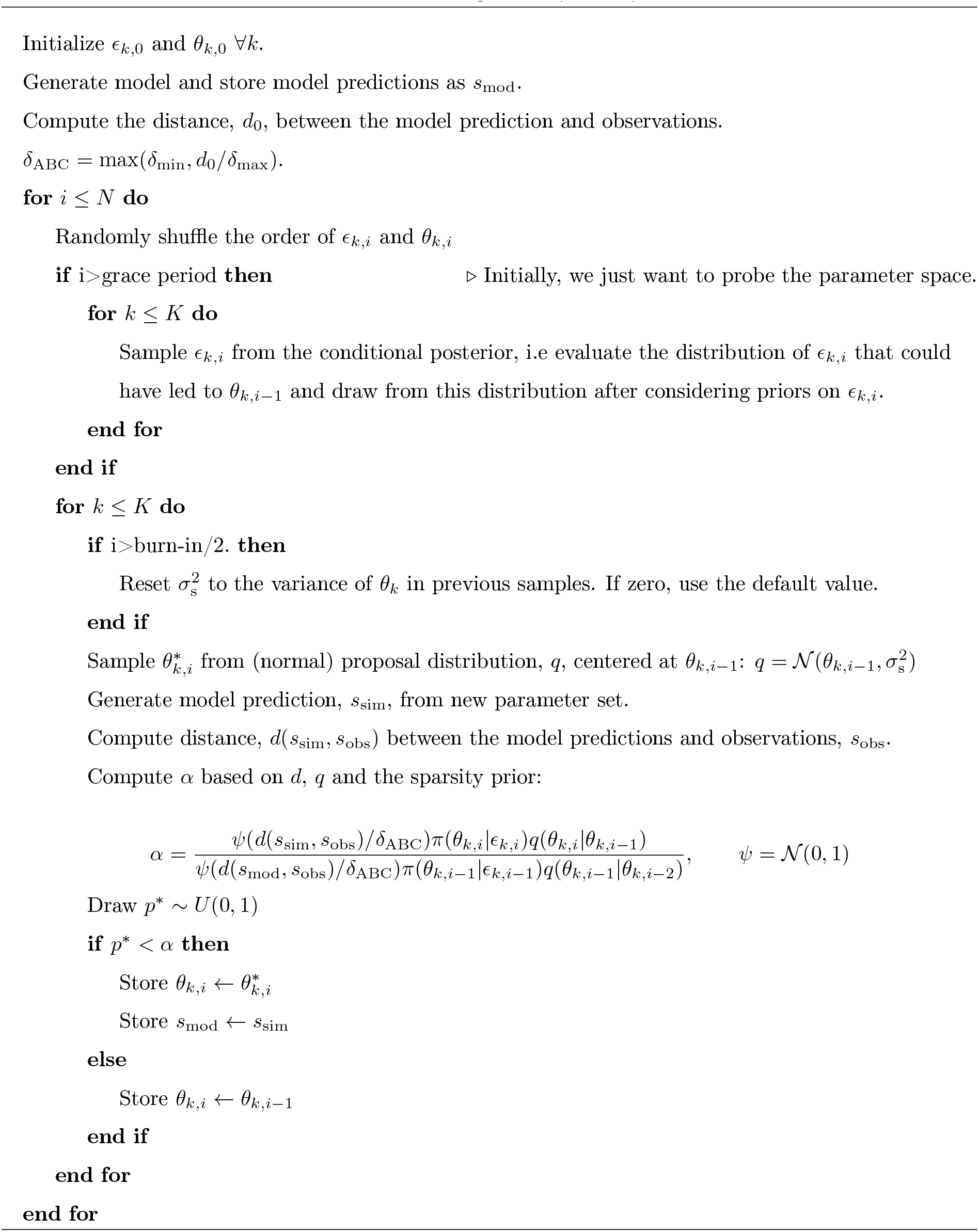

The algorithm might be called several times during the burn-in phase to adjust *δ*_ABC_ iteratively to the value set by the user *(δ*_min_). The grace period is only included during the first of these iterations. During the grace period, *ϵ*_*k,i*_ is set to 1 for all k.

## B Supplementary methods

### B.1 Linear models, settings

We use the diabetes dataset from Efron et al. provided through Scikit-learn [46].

We fitted the data set from Efron et al. using four different values of *δ*_ABC_. In all four cases, we imposed truncated Laplace priors, restricting parameters to values between -1,000 and 1,000.

We constructed 100 synthetic data sets. To create a synthetic data set, we randomly select between 1 and 6 of the 10 parameters and let these parameters take random values between 50 and 900. We then compute the resulting model response using our linear model. Finally, to ensure a realistic behaviour of the synthetic data, we add 5 per cent Gaussian noise to each data point, i.e. each data point *x*_*j*_ in the original synthetic data set is substituted by 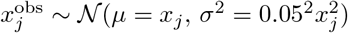.

To map the impact of *δ*_ABC_, we repeat the inference for each of the 100 data sets using 66 different values of *δ*_ABC_ between 0.00125 and 0.05. We thus perform 6,600 separate inference tasks using the same truncated Laplace priors as in the case of the real-world data. For this analysis, we consider any parameter that takes a value lower than 10 as discarded since the relevant parameter range stretches from 50 to 900.

### B.2 Adaptation, ODEs

The considered adaptive regulatory networks [see also 35] are described by a system of three ordinary differential equations (ODEs):

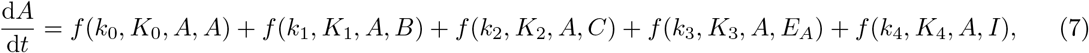

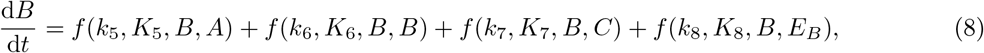

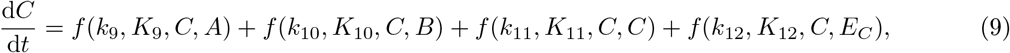

where *A, B*, and *C* denote the three genes/enzymes in the regulatory network, while *E*_*A*_, *E*_*B*_, and *E*_*C*_ are other activating/deactivating enzymes that might affect *A, B*, or *C*, respectively. *I* denotes the input signal, and *k*_0_-*k*_12_ and *K*_0_-*K*_12_ are the 26 free parameters that we aim to fit. Finally, *f* denotes the Hill function

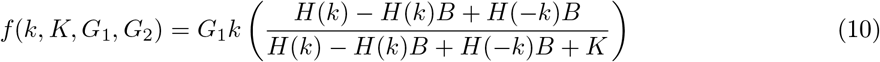

where *H*(*k*) denotes the Heaviside function, i.e. the sign of *k* determines whether the link leads to repression or activation.

### B.3 Adaptation, settings

For all parameters in equations (7-9) in Appendix B.2, we use uniform priors. We allow *k*_0_-*k*_12_ (with the exception of *k*_4_) to take values between -10 and 10, i.e. all links might either be repressive or activating. We allow *K*_0_ − *K*_12_ as well as *k*_4_ to vary between 0.1 and 10. The reason we single out *k*_4_ is that we only consider networks in which the input activates the first node (*A*).

We note that there must be both positive and negative terms on the right-hand side of Equations 7-9 since the concentration of A, B, and C will otherwise diverge. Thus, we outright discard all models for which the sum over the negative link strengths or the sum over positive link strengths falls below a protein-specific threshold, i.e. the Markov chain does not pass through such points. For A and C, we set the threshold to 10^*−*3^, while we set the threshold to 10^*−*4^ when dealing with B. These thresholds are somewhat arbitrary. Changing them would further limit or expand the parameter space that we explore. We also note that B is treated differently since it’s a buffer node/proportioner: According to Ma et al., the presence of such nodes is essential for achieving adaptation, making us more reluctant to discard links to B.

We set up one hundred runs starting each run from a fully connected network with uniform link strengths of 0.1. The one hundred runs are subdivided into groups of twenty, each group using a different value of *δ*_ABC_: 0.5, 0.2, 0.1, 0.05, and 0.029. The last value (1/35) in this list is determined by the distance measure of the initial conditions and is, hence, the lowest possible value for our set-up (we set *δ*_max_ = 35, cf. Appendix A). For each run, we create a single Markov chain with 10,000 samples.

After each run, we extract the last 4,000 models in the Markov chain and set all link strengths that fall below a certain threshold to zero. We vary this threshold, increasing it from 10^*−*3^ to 10^*−*2^ and, finally, to 10^*−*1^. We then recompute *P* and *S* for each of the models — as both *P* and *S* will change when setting some parameters to zero. We then identify the adaptive networks, i.e. networks with *S* > 1 and *P* > 10.

Among the 31 successful topologies, *δ*_ABC_ was set to 0.029, 0.05 and 0.1 in 7, 18 and 6 cases, respectively, while no adaptive systems were found when choosing higher values of *δ*_ABC_.

### B.4 Stochastic regulatory genetic networks, SDEs

The considered stochastic regulatory genetic networks are represented by a system of eight stochastic differential equations (SDEs):

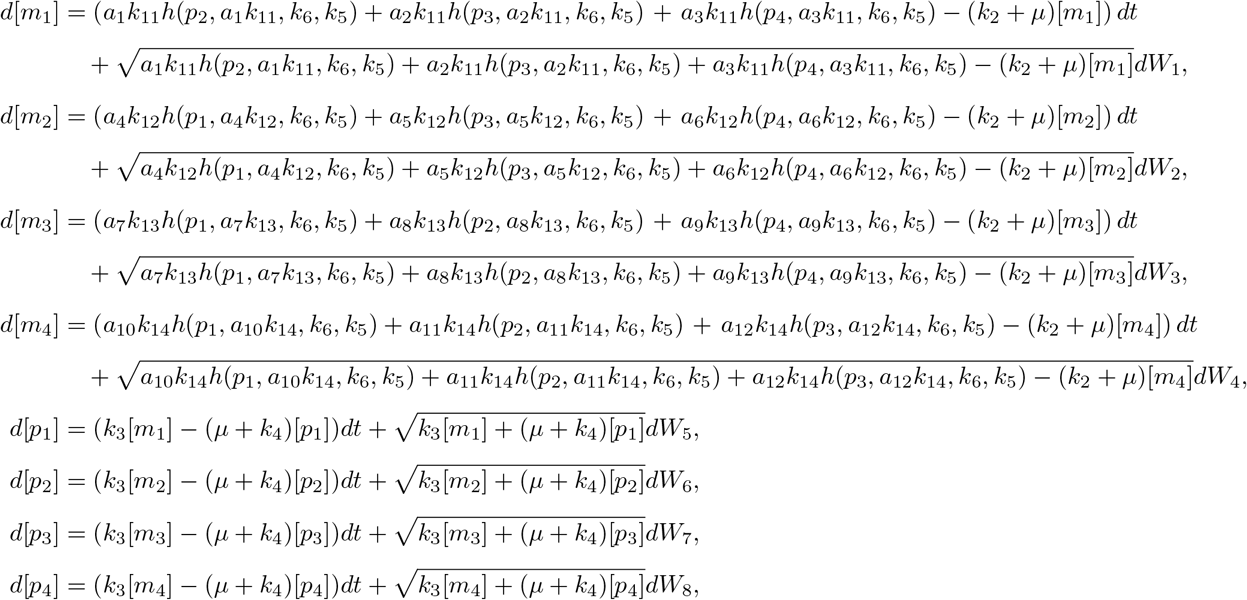

where [*m*_1_] to [*m*_4_] represent mRNA concentrations for each gene, while [*p*_1_] to [*p*_4_] represent protein concentrations for each gene. *W*_1_ to *W*_8_ are independent Wiener processes representing stochastic noise in the reactions. Finally, activation or repression interactions are represented by

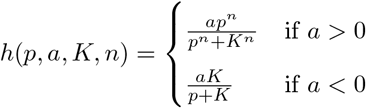

where *k*_11_, *k*_12_, *k*_13_, and *k*_14_ represent the basal transcription rates for each of the four genes, respectively. Moreover, *k*_2_ represents the rate of mRNA decay, *k*_3_ represents the translation rate, and *k*_4_ is the rate of protein decay. Cell dilution is modelled to occur as a result of exponential cell growth and is characterised by the rate *µ*, while *k*_5_ is a hill coefficient, and *k*_6_ is a dissociation constant. Sparse genetic networks can be created by setting the link strength parameters, *a*_1_ to *a*_12_, to zero.

We note that the impact of the strengths of the different regulatory links on mean gene expression can be mimicked by changing the base rates of gene expression for the four genes (i.e. *k*_11_ − *k*_14_) — to this end, the base rates and the link strengths are degenerate. In real-world applications, *k*_11_ − *k*_14_ are unknown and must hence be inferred alongside the strength of the different links between the genes. We thus require SLInG to infer *k*_11_ − *k*_14_ as well as 12 parameters, one associated with the possible links between the genes, i.e. we have 16 free parameters in total.

Because we do not impose any sparsity priors on *k*_11_ − *k*_14_ (cf. Appendix B.5), the sampler increases *k*_11_ −*k*_14_ above the ground truth to reduce the link strengths and potentially discard links. This behaviour is due to the mentioned degeneracy between the impact of *k*_11_ − *k*_14_ and the link strengths on the mean gene expression. SLInG thus points to the existence of links based on the contribution of the covariance matrix to the distance measure rather than based on the mean gene expression.

### B.5 Stochastic regulatory genetic networks, settings

For the 12 link strengths, we impose truncated Laplace priors, constraining them to take values between -20 and 20: Negative link strengths imply that the corresponding link is repressive, while positive links lead to activation. For *k*_11_ − *k*_14_, on the other hand, we do not impose sparsity priors but merely impose uniform priors on their logarithm from -2 to 2.

We created 100 synthetic data sets based on random 4-node gene-regulatory networks. For each data set, we randomly selected the total number of existing (non-zero) links, the number of repressive and adaptive links amongst the non-zero links, and the strength of each of the non-zero links. Thus, each synthetic data set stems from a network with between 3 and 5 non-zero links that were each randomly set to be repressive or activating — both choices were equally likely. The absolute strength of the links all take values between 3.0 and 10.0, i.e. safely within the prior of the sampler. Because the detection of links relies on the contribution of the covariance matrix to the distance measure, we further required that all synthetic data sets show significant correlations between the gene expressions. We thus only adopted a randomly chosen parameter set into our library of synthetic data sets if at least six out of the twelve off-diagonal elements in the *correlation* matrix of the resulting synthetic data were larger than 0.3. Otherwise, we tried another parameter set. In other words, the 100 synthetic data sets are selected using trial and error to fulfil our condition on the correlation matrix. For all runs, *k*_11_ − *k*_14_ are set to 0.40, 0.37, 0.34 and 0.30, respectively.

In the paper, we increase the value of *δ*_ABC_ to make the sampler less sensitive to the stochasticity of the simulations. More specifically, we found that a value of 0.15 allows our sampler to efficiently explore the parameter space while still capturing the essential information in the data for most parameter sets. In some cases, we found that the chain still seemed to get stuck. If we were dealing with real-world data, we would try different values of *δ*_ABC_ to fit the data. However, rather than choosing a larger value for *δ*_ABC_ for these cases, we simply discarded the synthetic data set, created a new one, and repeated the analysis. This choice was merely made to ensure a uniform value of *δ*_ABC_ across the samples.

To infer the parameter values that underlie the synthetic data, we initialised each run, setting all parameters to zero, including the logarithm of *k*_11_ − *k*_14_. For each of the 100 runs, we computed 5,500 samples — each sample involves 16 simulations, as we are exploring a 16-dimensional parameter space. The initial 1,500 samples functioned as a burn-in phase, which allowed us to adjust *δ*_ABC_ iteratively. For the analysis of each run, we focus on the best-fitting model amongst the 4,000 samples that were left after excluding this burn-in phase. As a first quality check, we determine whether the best-fitting parameter set produces gene expressions that lie sufficiently close to the data. This comparison is performed in three steps: First, we bootstrap the observations, i.e. the synthetic data, and compute the distance of each bootstrapped data set to the observations. Secondly, we compute the gene expression of the best-fitting parameter set after discarding any links with strengths below 10^*−*1^. This cut-off was set based on the following considerations: We only want to keep the most prominent links. However, while no data set is constructed with a link strength lower than 3.0, the inference will yield lower values due to the aforementioned degeneracy between the link strengths and *k*_11_ − *k*_14_.

Thirdly, we bootstrap the results for the best-fitting model, computing the distance between each bootstrapped set of predictions and the observations. If the resulting distribution of distances overlaps with even one sample from the distribution of the distance between the bootstrapped data and the observations from the first step of our comparison discussed above, we accept the fit. Otherwise, we discard the fit. Were we forced to discard a fit when applying the algorithm to real-world data, we could try to improve the fit by, e.g., adjusting initial conditions. However, as mentioned in the paper, we merely note that the algorithm passed our quality check out of the box in 56 per cent of the cases — this success rate is not significantly affected by the cut-off in link strength for determining the number of non-zero links.

### B.6 Other algorithms for reverse engineering signalling networks

GENIE3, an acronym for GEne Network Inference with Ensemble of trees, was the best performer at the DREAM4/DREAM5 network challenges [38], and though developed more than a decade ago, it still remains a favoured method for this kind of inference [22, 26]. The algorithm uses an ensemble of decision trees to model the expression of a gene as a function of the expression of other genes. Its primary assumption is that the expression of a gene is the combined effect of the expression of other genes, which is captured by the tree structure. This leads to a directed network where the genes are connected based on their inferred influence on each other.

GRNBoost2 is a more recently developed algorithm for gene regulatory network inference and is based on the GENIE3 architecture [41]. Despite sharing the same aim to uncover complex gene interactions, the two methods employ different machine-learning strategies. GRNBoost2 employs gradient boosting machines, an ensemble learning method that refines new models iteratively to provide a more precise approximation of the response variable [16]. In the context of GRNBoost2, the response variable corresponds to the expression level of a target gene. With each iteration, GRNBoost2 works to minimise the residuals between the predicted and actual expression levels of a target gene, successively refining its model. It constructs a network by identifying the relationships between a transcription factor and a target gene that maximise shared mutual information, taking into account the effects of all other transcription factors. Although GRNBoost2 operates under a different learning method, its ultimate goal aligns with that of GENIE3: It aims to generate an accurate, directed network that represents gene regulatory relationships.

The practical implementation of GENIE3 and GRNBoost2 in large-scale network inference tasks can be computationally demanding. To overcome this issue, we made use of the Arboreto library, a highly efficient Python framework that facilitates the application of both these algorithms [41]. Arboreto has been optimised for scalability and performance. By leveraging Dask, a flexible parallel computing library for analytic computing [61], Arboreto is able to distribute and parallelise the execution of the GRN inference tasks over multiple cores or nodes. It also employs smart scheduling to prioritise the most crucial tasks, thereby maximising computational efficiency.

The application of GENIE3 and GRNBoost2 results in an ordered list of interactions ranked in terms of their inferred ‘importance’ or regulatory significance within the gene network. However, as these are stochastic methods, they introduce an element of randomness in their process, potentially leading to slight variations in outcomes across different runs. To mitigate this stochastic nature and provide robustness to our results, we conducted 100 repeats of the network inference process and aggregated the results by taking an average across all runs. We then used an Elbow-method to estimate the number of interactions from the full list. Finally, we used the ‘ppcor’ package in R to compute the directionality of the interactions [25] and produce estimates of signed, directed networks that we can compare with SLInG.

## C Supplementary figures

**Figure S1:**
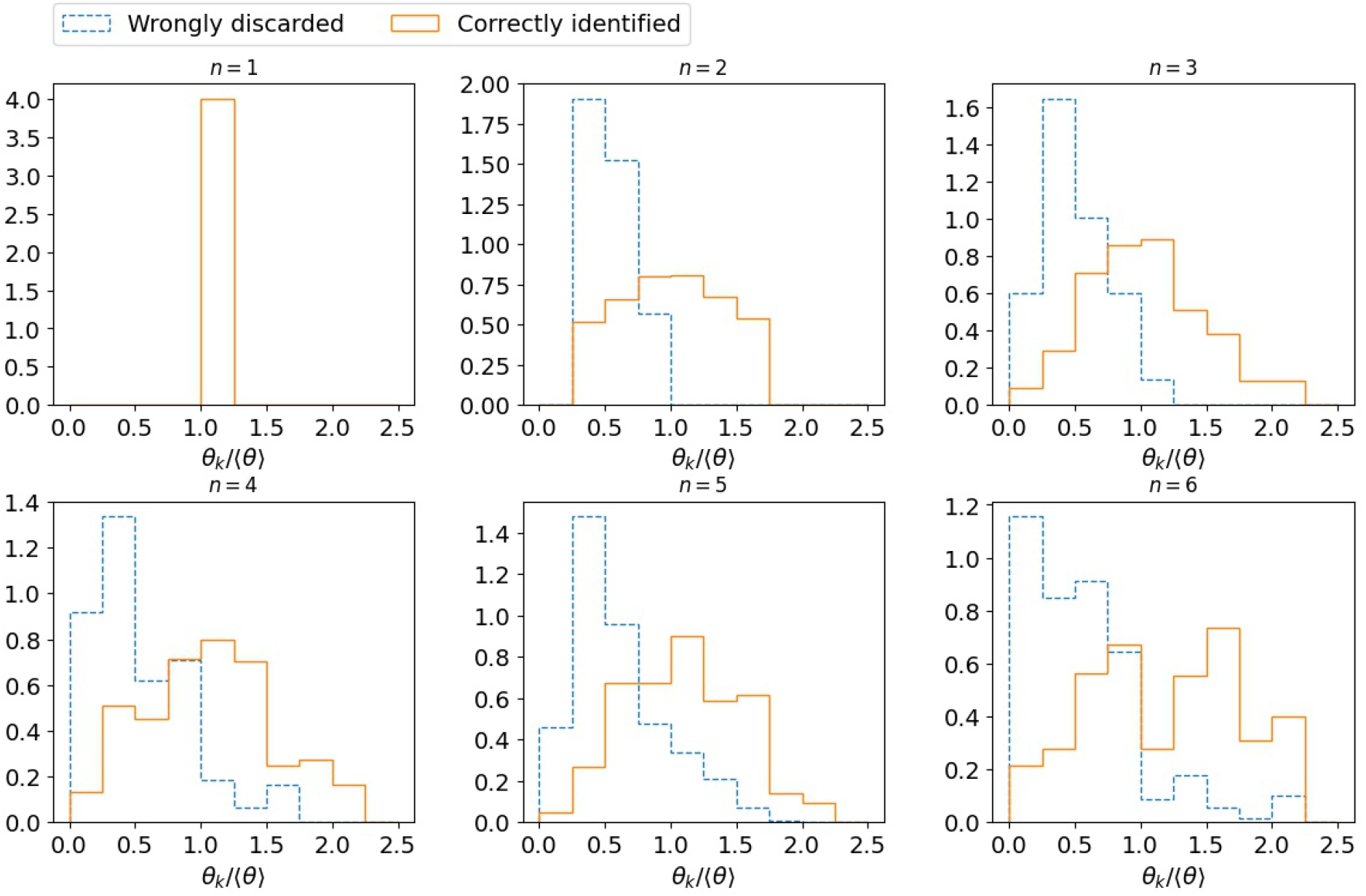
Parameter values across the different number of parameters (*n*) employed to produce the synthetic data for our linear model in Section 3.1. The plot compares the true values of the parameters that were correctly identified with the values of those that were not found by SLInG. The plot includes the same 6600 simulations presented in Fig 3 in the main text. The parameters that are not correctly identified by the algorithm tend to have the weakest impact on the response variable. They thus take values that are lower than the mean parameter values across the n true parameters (⟨ *θ*⟩). In contrast, SLInG correctly points to the parameters with the highest explanatory power.

**Figure S2:**
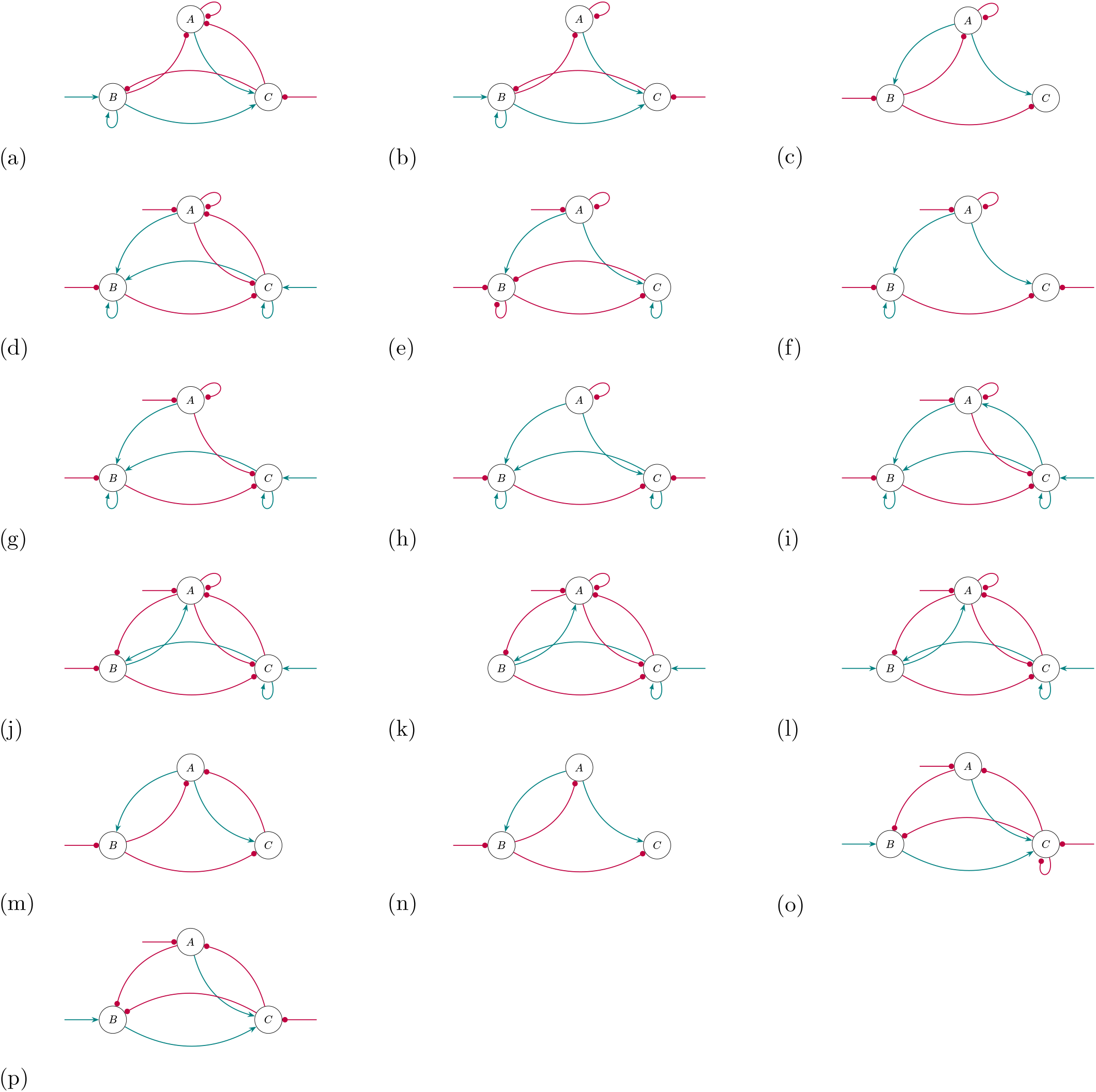
16 of the 31 adaptive regulatory networks found by the sampler.

**Figure S3:**
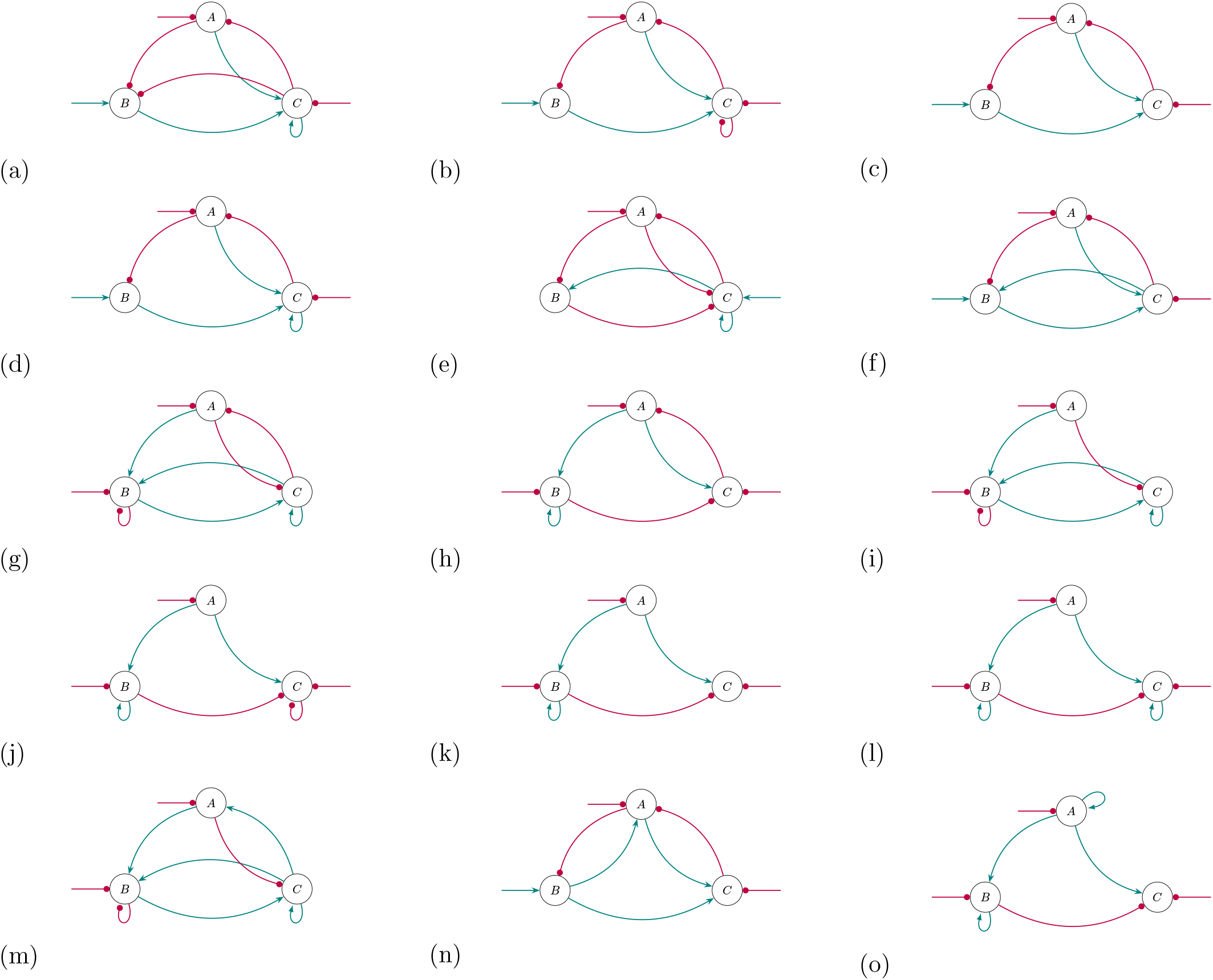
15 of the 31 adaptive regulatory networks found by the sampler.

**Figure S4:**
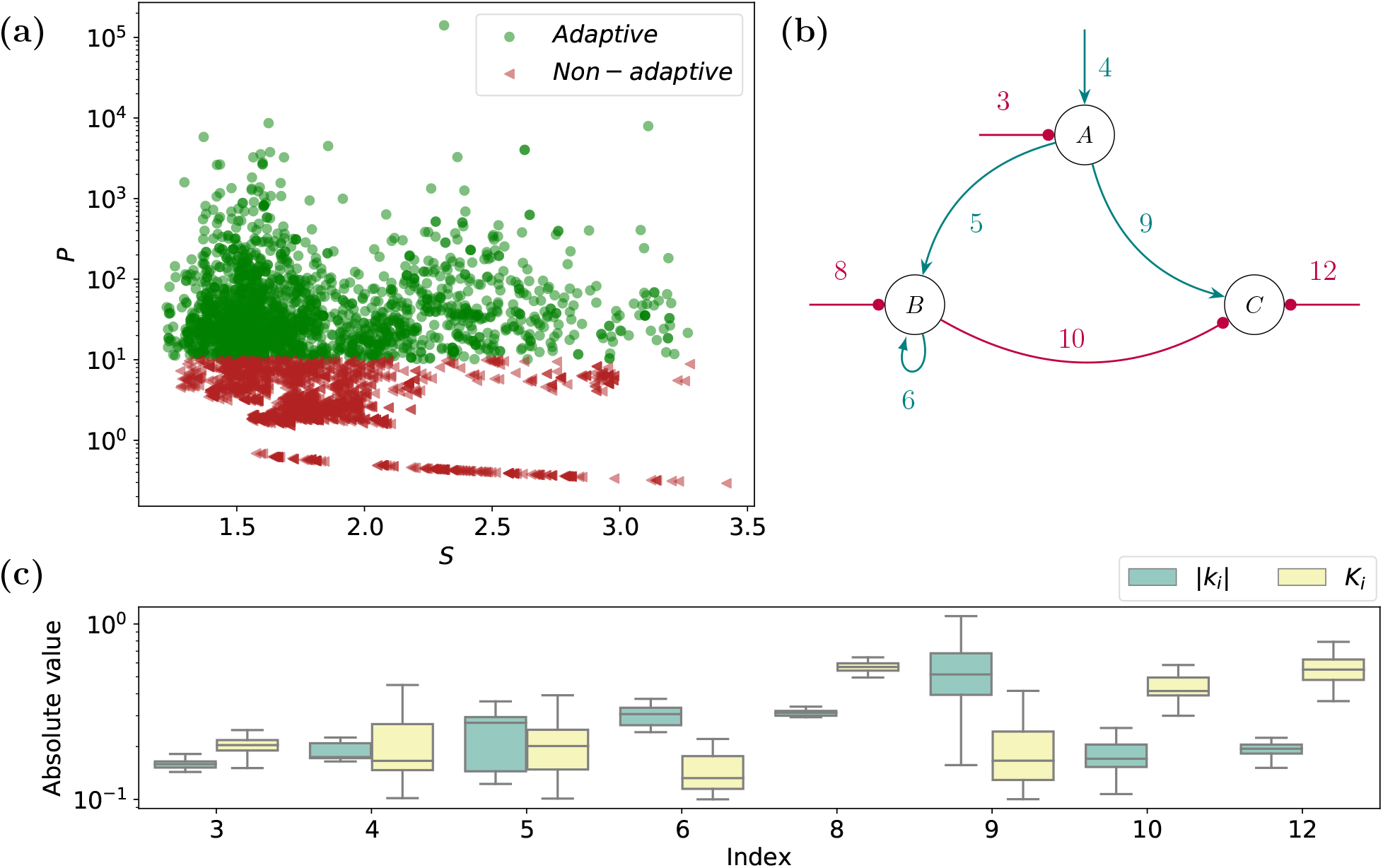
The impact of changing parameter values on *P* and *S* for the topology in panel (l) of Fig S3. (a) *P* as a function of *S* for all samples with this topology among the 4,000 last samples of the Markov chain. (b) The topology of the signalling network with the indices of the corresponding terms from Equations (7)-(9). (c) The parameter values *that lead to adaptation* within the samples. The index (*i*) corresponds to the link number in panel b. As can be seen from the figure, not all parameter combinations lead to adaptation, although adaptation is preserved for a range of parameter values.

**Figure S5:**
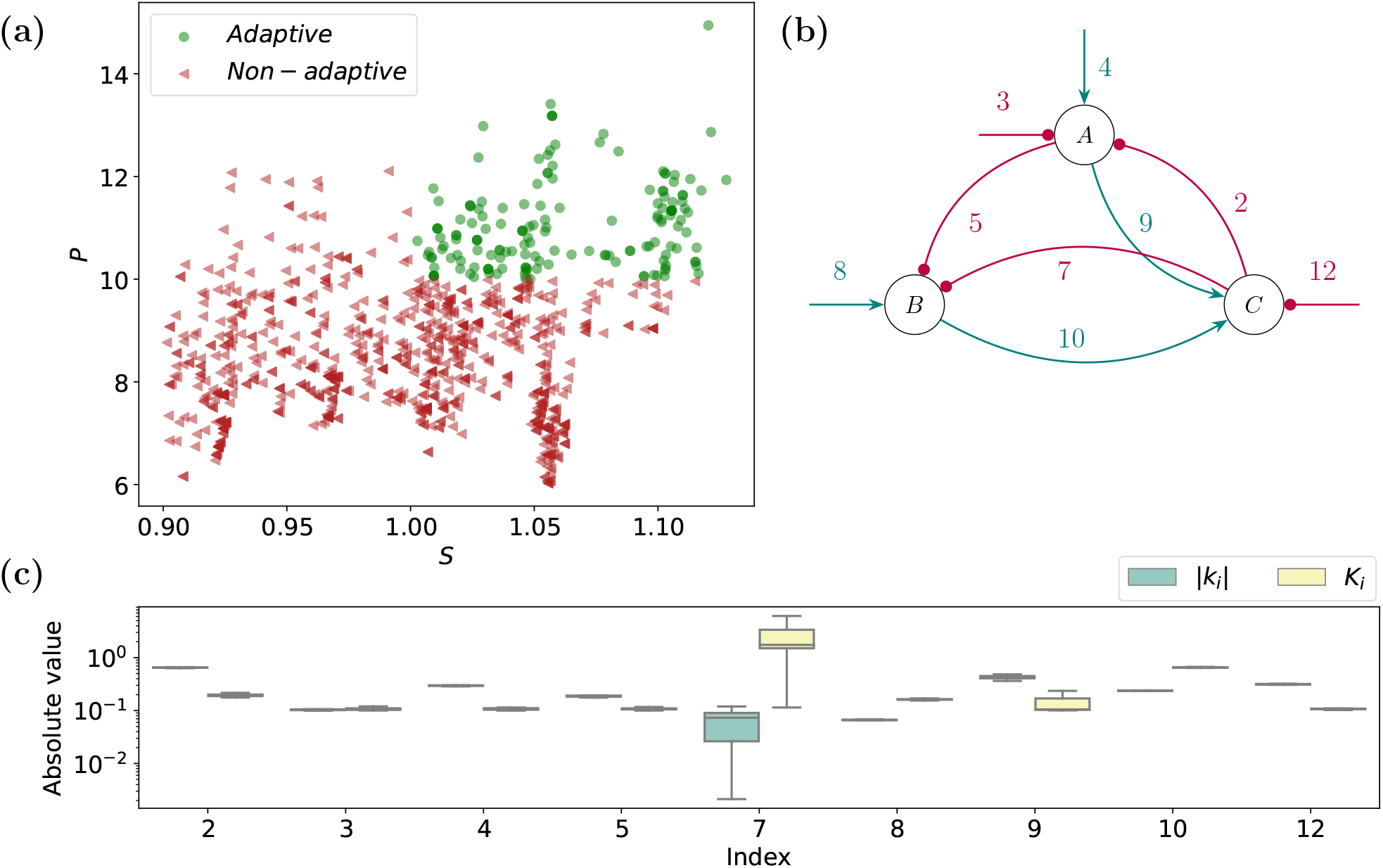
Pendant for Fig S4 for panel (p) in Fig S3. Again, not all parameter combinations lead to adaptation. Indeed, only a very narrow range of samples leads to adaptation.

## References

[1] V. Aguirre Børsen-Koch, J. L. Rørsted, A. B. Justesen, A. Stokholm, K. Verma, M. L. Winther, E. Knudstrup, K. B. Nielsen, C. Sahlholdt, J. R. Larsen, S. Cassisi, A. M. Serenelli, L. Casagrande, J. Christensen-Dalsgaard, G. R. Davies, J. W. Ferguson, M. N. Lund, A. Weiss, and T. R. White. The BAyesian STellar algorithm (BASTA): a fitting tool for stellar studies, asteroseismology, exoplanets, and Galactic archaeology. MNRAS, 509(3):4344–4364, 2022.

[2] Amani A. Alahmadi, Jennifer A. Flegg, Davis G. Cochrane, and Jonathan M. Drovandi, Christopher C. Keith. A comparison of approximate versus exact techniques for bayesian parameter inference in nonlinear ordinary differential equation models. Royal Society open science, 2020.

[3] Rahim Alhamzawi and Haithem Taha Mohammad Ali. A new gibbs sampler for bayesian lasso. Communications in Statistics - Simulation and Computation, 49(7):1855–1871, 2020.

[4] Małgorzata Bogdan, Ewout van den Berg, Chiara Sabatti, Weijie Su, and Emmanuel J. Candès. Slope - adaptive variable selection via convex optimization. The annals of applied statistics, 9:1103–1140, 2015.

[5] Gert-Jan Both, Subham Choudhury, Pierre Sens, and Remy Kusters. Deepmod: Deep learning for model discovery in noisy data. Journal of Computational Physics, 428:109985, 2021.

[6] Steven L. Brunton, Joshua L. Proctor, and J. Nathan Kutz. Discovering governing equations from data by sparse identification of nonlinear dynamical systems. Proceedings of the National Academy of Science, 113(15):3932–3937, 2016.

[7] Maria Carilli, Gennady Gorin, Yongin Choi, Tara Chari, and Lior Pachter. Biophysical modeling with variational autoencoders for bimodal, single-cell rna sequencing data, 2023.

[8] Xiaohui Chen, Z. Jane Wang, and Martin J. McKeown. A bayesian lasso via reversible-jump mcmc. Signal Processing, 91(8):1920–1932, 2011.

[9] Bradley Efron, Trevor Hastie, Iain Johnstone, and Robert Tibshirani. Least angle regression. The Annals of Statistics, 32(2):407–451, 2004.

[10] Robert G. Endres and Ned S. Wingreen. Precise adaptation in bacterial chemotaxis through “assistance neighborhoods”. Proceedings of the National Academy of Sciences of the United States of America, 103:13040–4, 2006.

[11] Song Feng, Julien F. Ollivier, Peter S. Swain, and Orkun S. Soyer. Biojazz: in silico evolution of cellular networks with unbounded complexity using rule-based modeling. Nucleic Acids Research, 43(19):e123–e123, 06 2015.

[12] Fabio Feser and Marina Evangelou. Sparse-group SLOPE: adaptive bi-level selection with FDR-control. arXiv e-prints, page arXiv:2305.09467, 2023.

[13] Paul François and Vincent Hakim. Design of genetic networks with specified functions by evolution in silico. Proceedings of the National Academy of Sciences of the United States of America, 101:580–585, 2004.

[14] Paul François and Eric D Siggia. Predicting embryonic patterning using mutual entropy fitness and in silico evolution. Development (Cambridge, England), 137(14):2385—2395, July 2010.

[15] David T. Frazier, Christian P. Robert, and Judith Rousseau. Model misspecification in approximate bayesian computation: consequences and diagnostics. Journal of the Royal Statistical Society Series B-Statistical Methodology, 82(2):421–444, April 2020.

[16] Jerome H. Friedman. Greedy function approximation: A gradient boosting machine. The Annals of Statistics, 29(5):1189–1232, 2001.

[17] R. Fuentes, R. Nayek, P. Gardner, N. Dervilis, T. Rogers, K. Worden, and E.J. Cross. Equation discovery for nonlinear dynamical systems: A bayesian viewpoint. Mechanical Systems and Signal Processing, 154:107528, 2021.

[18] Daniel T. Gillesple. Exact stochastic simulation of coupled chemical reactions. J. Phys. Chem., 81:2340–2361, 1977.

[19] Lucy Ham, Lucy Ham, Marcel Jackson, and Michael Ph Stumpf. Pathway dynamics can delineate the sources of transcriptional noise in gene expression. eLife, 10:e69324, October 2021.

[20] Clinton H Hansen, Robert G Endres, and Ned S Wingreen. Chemotaxis in escherichia coli: a molecular model for robust precise adaptation. PLoS computational biology, 4(1):e1, January 2008.

[21] S. M. Hirsh, D. A. Barajas-Solano, and J. N. Kutz. Sparsifying priors for bayesian uncertainty quantification in model discovery. Royal Society Open Science, 9, 2022.

[22] Vân Anh Huynh-Thu, Alexandre Irrthum, Louis Wehenkel, and Pierre Geurts. Inferring regulatory networks from expression data using tree-based methods. PloS one, 5(9):e12776, 2010.

[23] Gareth James, Daniela Witten, Trevor Hastie, and Robert Tibshirani. An Introduction to Statistical Learning: with Applications in R. Springer, 2013.

[24] A. C. S. Jørgensen, A. Ghosh, M. Sturrock, and V. Shahrezaei. Efficient bayesian inference for stochastic agent-based models. PLoS computational biology, 18, 2022.

[25] Seongho Kim. ppcor: an r package for a fast calculation to semi-partial correlation coefficients. Communications for statistical applications and methods, 22(6):665, 2015.

[26] Ayoub Lasri, Vahid Shahrezaei, and Marc Sturrock. Benchmarking imputation methods for network inference using a novel method of synthetic scrna-seq data generation. BMC bioinformatics, 23(1):236, 2022.

[27] Ju-Sung Lee, Tatiana Filatova, Arika Ligmann-Zielinska, Behrooz Hassani-Mahmooei, Forrest Stonedahl, Iris Lorscheid, Alexey Voinov, J. Polhill, Zhanli Sun, and Dawn Parker. The complexities of agent-based modeling output analysis. Journal of Artificial Societies and Social Simulation, 18, 10 2015.

[28] Kyoungjae Lee, Jaeyong Lee, and Sarat C. Dass. Inference for differential equation models using relaxation via dynamical systems. Computational Statistics & Data Analysis, 127:116–134, 2018.

[29] Ismael Lemhadri, Feng Ruan, and Robert Tibshirani. Lassonet: Neural networks with feature sparsity. Proceedings of machine learning research, 130:10—18, April 2021.

[30] Maxime Lenormand, Franck Jabot, and Guillaume Deffuant. Adaptive approximate bayesian computation for complex models. Computational Statistics, 28, 11 2011.

[31] Melissa Lever, Hong-Sheng Lim, Philipp Kruger, John Nguyen, Nicola Trendel, Enas Abu-Shah, Philip Kumar Maini, Philip Anton van der Merwe, and Omer Dushek. Architecture of a minimal signaling pathway explains the t-cell response to a 1 million-fold variation in antigen affinity and dose. Proceedings of the National Academy of Sciences of the United States of America, 113, 2016.

[32] Jana Lipková, Panagiotis Angelikopoulos, Stephen Wu, Esther Alberts, Benedikt Wiestler, Christian Diehl, Christine Preibisch, Thomas Pyka, Stephanie E. Combs, Panagiotis Hadjidoukas, Koen Van Leemput, Petros Koumoutsakos, John Lowengrub, and Bjoern Menze. Personalized radiotherapy design for glioblastoma: Integrating mathematical tumor models, multimodal scans, and bayesian inference. IEEE Transactions on Medical Imaging, 38(8):1875–1884, 2019.

[33] T. Loman, Y. Ma, V. Ilin, S. Gowda, N. Korsbo, N. Yewale, C. V. Rackauckas, and S. A. Isaacson. Catalyst: Fast biochemical modeling with julia. bioRxiv, 2022.

[34] Jan-Matthis Lueckmann, Jan Boelts, David S. Greenberg, Pedro J. Gonçalves, and Jakob H. Macke. Benchmarking Simulation-Based Inference. arXiv e-prints, page arXiv:2101.04653, January 2021.

[35] Wenzhe Ma, Ala Trusina, Hana El-Samad, Wendell A Lim, and Chao Tang. Defining network topologies that can achieve biochemical adaptation. Cell, 138(4):760—773, August 2009.

[36] Benn Macdonald and Dirk Husmeier. Gradient matching methods for computational inference in mechanistic models for systems biology: A review and comparative analysis. Frontiers in Bioengineering and Biotechnology, 3(NOV):1–21, 2015.

[37] Himel Mallick and Nengjun Yi. A new bayesian lasso. Statistics and its interface, 7:571–582, 2014.

[38] Daniel Marbach, James C Costello, Robert Küffner, Nicole M Vega, Robert J Prill, Diogo M Camacho, Kyle R Allison, Manolis Kellis, James J Collins, et al. Wisdom of crowds for robust gene network inference. Nature methods, 9(8):796–804, 2012.

[39] Jean-Michel Marin, Pierre Pudlo, Christian P. Robert, and Robin Ryder. Approximate bayesian computational methods. Statistics and Computing, 22:1167 –1180, 2012.

[40] Jameson Miller, Miles Parker, Robert B Bourret, and Morgan C Giddings. An agent-based model of signal transduction in bacterial chemotaxis. PloS one, 5(5):e9454, May 2010.

[41] Thomas Moerman, Sara Aibar Santos, Carmen Bravo González-Blas, Jaak Simm, Yves Moreau, Jan Aerts, and Stein Aerts. Grnboost2 and arboreto: efficient and scalable inference of gene regulatory networks. Bioinformatics, 35(12):2159–2161, 2019.

[42] R. Nayek, R. Fuentes, K. Worden, and E. J. Cross. On spike-and-slab priors for bayesian equation discovery of nonlinear dynamical systems via sparse linear regression. Mechanical Systems and Signal Processing, 161:107986, 2021.

[43] David Nott, Christopher Drovandi, and David Frazier. Bayesian inference for misspecified generative models. Annual Review of Statistics and Its Application, 11, 08 2023.

[44] R. B. O’Hara and M. J. Sillanpää. A review of bayesian variable selection methods: what, how and which. Bayesian Analysis, 4(1):85 –117, 2009.

[45] Trevor Park and George Casella. The bayesian lasso. Journal of the American Statistical Association, 103(482):681–686, 2008.

[46] F. Pedregosa, G. Varoquaux, A. Gramfort, V. Michel, B. Thirion, O. Grisel, M. Blondel, P. Pretten-hofer, R. Weiss, V. Dubourg, J. Vanderplas, A. Passos, D. Cournapeau, M. Brucher, M. Perrot, and E. Duchesnay. Scikit-learn: Machine learning in Python. Journal of Machine Learning Research, 12:2825–2830, 2011.

[47] William Pontius, Michael W Sneddon, and Thierry Emonet. Adaptation dynamics in densely clustered chemoreceptors. PLoS computational biology, 9(9):e1003230, 2013.

[48] Natalia Porqueres, Alan Heavens, Daniel Mortlock, Guilhem Lavaux, and T. Lucas Makinen. Field-level inference of cosmic shear with intrinsic alignments and baryons. arXiv e-prints, page arXiv:2304.04785, 2023.

[49] Xiaojie Qiu, Yan Zhang, Jorge D Martin-Rufino, Chen Weng, Shayan Hosseinzadeh, Dian Yang, Angela N Pogson, Marco Y Hein, Kyung Hoi Joseph Min, Li Wang, Emanuelle I Grody, Matthew J Shurtleff, Ruoshi Yuan, Song Xu, Yian Ma, Joseph M Replogle, Eric S Lander, Spyros Darmanis, Ivet Bahar, Vijay G Sankaran, Jianhua Xing, and Jonathan S Weissman. Mapping transcriptomic vector fields of single cells. Cell, 185(4):690—711.e45, February 2022.

[50] J Reisert and HR Matthews. Response properties of isolated mouse olfactory receptor cells. The Journal of physiology, 530(Pt 1):113—122, January 2001.

[51] Ben M. Rendle, Gaël Buldgen, Andrea Miglio, Daniel Reese, Arlette Noels, Guy R. Davies, Tiago L. Campante, William J. Chaplin, Mikkel N. Lund, James S. Kuszlewicz, Laura J. A. Scott, Richard Scuflaire, Warrick H. Ball, Jiri Smetana, and Benard Nsamba. AIMS - a new tool for stellar parameter determinations using asteroseismic constraints. MNRAS, 484(1):771–786, 2019.

[52] Samuel H. Rudy, Steven L. Brunton, Joshua L. Proctor, and J. Nathan Kutz. Data-driven discovery of partial differential equations. Science Advances, 3(4):e1602614, 2017.

[53] Natalie S Scholes, David Schnoerr, Mark Isalan, and Michael P H Stumpf. A comprehensive network atlas reveals that turing patterns are common but not robust. Cell systems, 9(5):515—517, November 2019.

[54] Vahid Shahrezaei and Peter S Swain. Analytical distributions for stochastic gene expression. Proceedings of the National Academy of Sciences of the United States of America, 105, 2008.

[55] Jingxiang Shen, Feng Liu, Yuhai Tu, and Chao Tang. Finding gene network topologies for given biological function with recurrent neural network. Nature Communications, 12:3125, 2021.

[56] Wenjia Shi, Wenzhe Ma, Liyang Xiong, Mingyue Zhang, and Chao Tang. Adaptation with transcriptional regulation. Scientific reports, 7, 2017.

[57] Michael P.H. Stumpf. Inferring better gene regulation networks from single-cell data. Current Opinion in Systems Biology, 27:100342, 2021.

[58] Mikael Sunnåker, Alberto Giovanni Busetto, Elina Numminen, Jukka Corander, Matthieu Foll, and Christophe Dessimoz. Approximate bayesian computation. PLOS Computational Biology, 9:e1002803, 01 2013.

[59] Amanda Swan, Thomas Hillen, John C Bowman, and Albert D Murtha. A patient-specific anisotropic diffusion model for brain tumour spread. Bulletin of mathematical biology, 80(5):1259—1291, May 2018.

[60] Evgeny Tankhilevich, Jonathan Ish-Horowicz, Tara Hameed, Elisabeth Roesch, Istvan Kleijn, Michael P H Stumpf, and Fei He. Gpabc: a julia package for approximate bayesian computation with gaussian process emulation. Bioinformatics (Oxford, England), 36(10):3286—3287, May 2020.

[61] Dask Development Team. Dask: Library for dynamic task scheduling, 2016.

[62] Robert Tibshirani. Regression shrinkage and selection via the lasso. Journal of the Royal Statistical Society. Series B (Methodological), 58(1):267–288, 1996.

[63] Tina Toni, David Welch, Natalja Strelkowa, Andreas Ipsen, and Michael P H Stumpf. Approximate bayesian computation scheme for parameter inference and model selection in dynamical systems. Journal of the Royal Society, Interface, 6(31):187—202, February 2009.

[64] Brandon M. Turner and Trisha Van Zandt. Hierarchical Approximate Bayesian Computation. Psychometrika, 79(2):185–209, 2014.

[65] David J. Warne, Ruth E. Baker, and Matthew J. Simpson. Rapid bayesian inference for expensive stochastic models. Journal of Computational and Graphical Statistics, 31(2):512–528, 2022.

[66] Christoph Zechner, Jakob Ruess, Peter Krenn, Serge Pelet, Matthias Peter, John Lygeros, and Heinz Koeppl. Moment-based inference predicts bimodality in transient gene expression. Proceedings of the National Academy of Sciences of the United States of America, 109(21):8340—8345, May 2012.

[67] Hui Zou. The adaptive lasso and its oracle properties. Journal of the American Statistical Association, 101(476):1418–1429, 2006.

